# A new concept in antiviral drug design yields a potent influenza inhibitor

**DOI:** 10.1101/2022.11.08.515737

**Authors:** Charley Mackenzie-Kludas, Wen-Yang Wu, Betty Jin, Jennifer L. McKimm-Breschkin, Ee Ling Seah, Paul Jones, Emily Fairmaid, Celeste M. K. Tai, Ding Yuan Oh, Peter Jenkins, Aeron C. Hurt, Lorena E. Brown

**Author notes:** These authors contributed equally: Charley Mackenzie-Kludas, Wen-Yang Wu.

## Abstract

Contemporary antiviral development, whether by rational drug design or forward pharmacology, primarily strives to produce ‘lock and key’ inhibitors. While the technology to identify druggable targets and create compounds to bind them has improved dramatically over the last century it has always been constrained by the finite availability of suitable binding sites that antiviral compounds can occupy. Here we present a new approach to drug design that utilizes compounds devised to alter the microenvironment of the virion surface making it incompatible with virus entry and illustrate this strategy with inhibitors of influenza virus. We show that compounds that produce a proton-rich mantle above the virion surface induce a conformational change in the viral hemagglutinin (HA) rendering the virus unable to interact with cellular receptors and gain entry to the cell. The compounds show exceptional antiviral activity both *in vitro* and *in vivo* and protect against influenza illness in mice and ferrets after a single dose, either therapeutically or prophylactically. The work presented here lays the foundation for a brand-new category of inhibitors that could be engineered to counter many different viruses and potentially other pathogens.

---

Currently licensed antiviral drugs for influenza target the neuraminidase (NA) to prevent efficient release of virus from cells^1^, or inhibit the functions of the viral polymerase^2^. These drugs block steps in the replication cycle rather than preventing initial infection. Compounds that theoretically could stop viral entry by blocking the receptor-binding pocket on the cell-attachment protein, HA, have remained elusive, potentially due to the shallow nature of the pocket^3^ and variability of surrounding amino acids. During infection, binding of the HA surface glycoprotein to its receptor on respiratory epithelial cells triggers endocytosis of the virus. Within the endosome, an increase in acidification causes a low pH-induced conformational change to take place in the HA^4^ to allow exposure of a hydrophobic “fusion peptide” and the extension and insertion of a coiled-coil trimer, tipped with the hydrophobic domain, into the endosomal membrane^5,6^. This process culminates in fusion of the viral envelope with the endosomal membrane and the creation of a fusion pore through which the viral genome can escape for replication. The native form of the HA is metastable^5,7^ and as such the shift to the low energy fusogenic form is irreversible^8^. The low pH conformation of the HA can no longer engage with cellular receptors^9^.

Here we validate our drug design concept using two compounds, MD185 (Fig. 1a) and MD345 (Fig.1b). These compounds were devised to induce an acidic milieu at the surface of free virions that mimics the low pH environment found inside the endosome, thus triggering premature initiation of the HA into its fusogenic conformation (Fig. 1c). MD185 performs this function via four aminomethanesulfonic acid groups connected in pairs to a pyromellitic acid backbone by D-aspartic acid, while the effector domain of MD345 is derived from a single chain of seven cysteic acid repeats connected to a tricarboxylic acid backbone (Supplementary Methods). In both cases, the acidic domains are linked to a sialic acid derivative related to the NA inhibitor, zanamivir^10,11^. Zanamivir binds to and inhibits the activity of the second virion surface glycoprotein NA, the functions of which include mediating efficient release of nascent virus from infected cells by removing sialic acid from virus receptors in the region of virus budding^1^. We have chosen a zanamivir-like compound as a virus-docking device because of its high barrier to the emergence of resistance^12^ and use the dimeric form which has been shown to have prolonged effectiveness *in vivo* compared to monomeric zanamivir ^13^.

**Figure 1.**
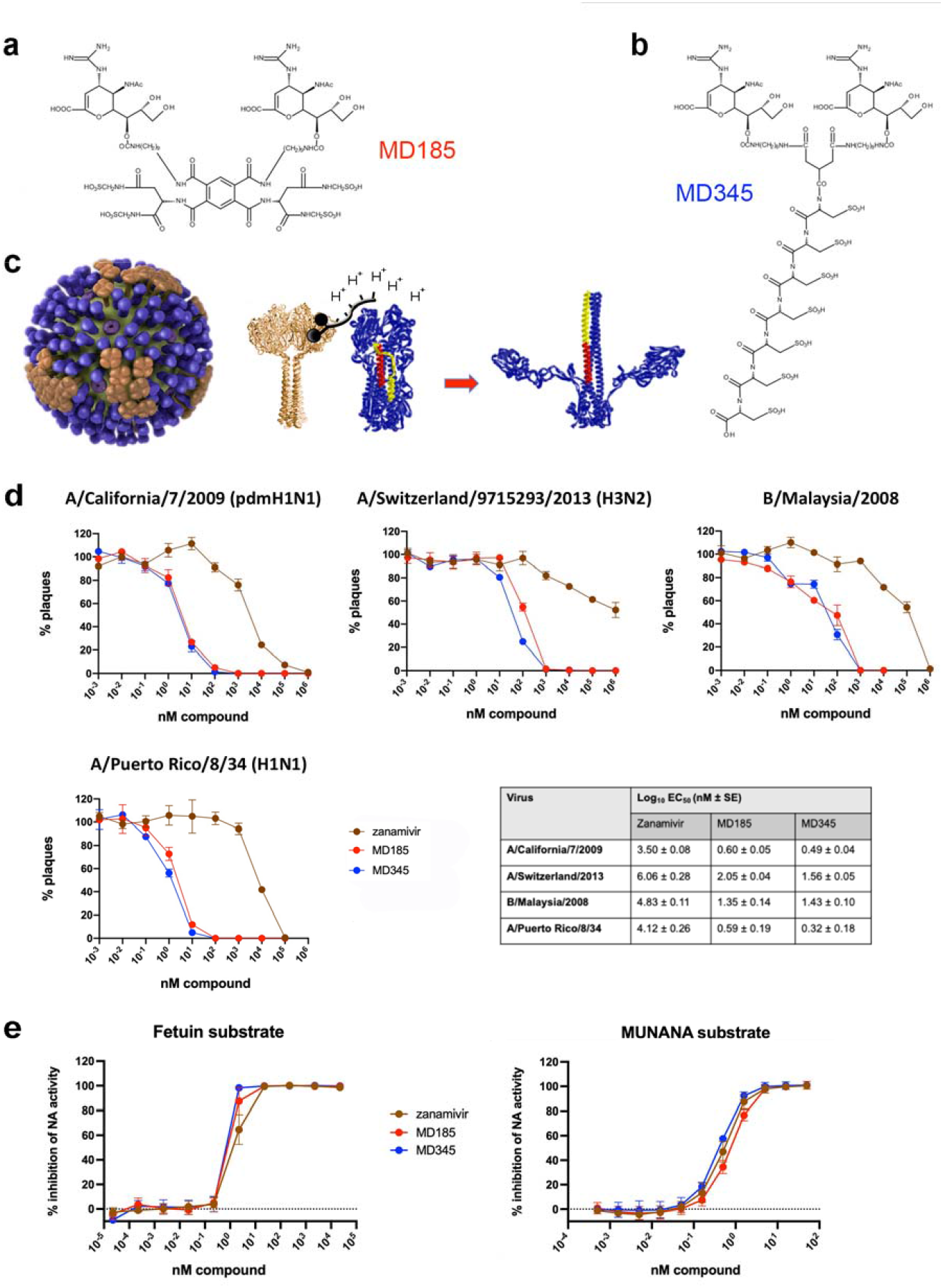
Structure, proposed action and *in vitro* activity of compounds. **a, b** The modular design of the compounds, which includes an NA-binding anchor, a structural backbone, and an acidic effector domain. For MD185 (**a**) the acidic domain consists of four aminomethanesulfonic acid groups connected to pyromellitic acid while MD345 (**b**) has a chain of seven cysteic acid repeats connected to a tricarboxcylic acid group. Both compounds are anchored to the virus by a pair of zanamivir homologs connected to the backbone via carbon chains. The molecular weight is 1852 and 2201 for MD185 and MD345 respectively. **c**, Graphical representation of the proposed mechanism of action. The compounds target the virion surface (left) and anchor to NA surface glycoprotein (brown). The acidic effector domain induces a drop in local pH (middle) that affects the neighbouring HA molecules (blue) causing the irreversible change in the HA from the metastable to the low pH fusogenic form (right) which precludes virion attachment to cell receptors. The illustration incorporates images of influenza virus from the Centres for Disease Control and Prevention, Atlanta, GA, USA; NA head domain, Protein Data Bank (PDB) 4GZX and the HA neutral form 1HGF. The extended intermediate fusion form is a schematic. **d**, Plaque reduction of PR8 (120 PFU), A/California/7/2009 (100 PFU), A/Switzerland/9715293/2013 (100 PFU), and B/Malaysia/2008 (100 PFU) influenza viruses by MD185 (red), MD345 (blue), or zanamivir (brown). All compounds were preincubated for 1 hr with virus before addition to MDCK cells and assayed for plaque formation. Data is expressed as a percent of plaques obtained with PR8 infection in the absence of compound. The Table insert shows the log EC_50_ of the compounds against each virus with the standard error of 3 determinations. **e**, PR8 NA inhibitory activity of the compounds. Data is calculated as a percentage of NA activity relative to PR8 in the absence of inhibitor and represents the average of 3 independent assays, each performed in duplicate, using fetuin (left) or MUNANA (right) as substrate.

Both MD185 and MD345 were effective against a range of influenza strains when screened in CPE reduction assays (Supplementary Table 1). In a plaque reduction assay, where viruses were pre-treated with compounds (Fig 1d), MD185 and MD345 showed 800-31,000-fold greater activity than zanamivir against human and mouse-adapted H1N1 viruses, a recent H3N2 strain, and an influenza B virus.This dramatic decrease in plaque number (Supplementary Fig.1) is not expected to result from pre-treatment of virus with an NA inhibitor. Usually assays of NA inhibitors require incorporation of drug into the overlay medium of the assay and this results primarily in reduction in plaque size, due to inhibition of virus spread, as opposed to plaque number, which suggests neutralization. Never-the-less, it was necessary to establish whether we had just created a superior NA inhibitor due to additional steric hindrance of the NA active site by the acidic groups on the bivalent docking domain. To investigate this, we performed enzyme inhibition assays using both the large sialylated substrate fetuin and the small artificial substrate 2’-(4-methylumbelliferyl)-α-D-N-acetylneuraminic acid (MUNANA). As can be seen in assays using either substrate (Figure 1e), there was no large improvement in the inhibition of NA activity by MD185 and MD345 compared to zanamivir. This demonstrates that while the anchor domains of the compounds do inhibit NA enzymatic function, this activity does not explain the very large increase in potency over zanamivir observed in the plaque reduction assays, suggesting that the enhanced efficacy of the compounds is due to an alternate mechanism.

To probe the effects of the compounds on HA we first tested whether the low pH conformation had indeed been induced after exposure to MD185 and MD345. Monoclonal antibodies (MAbs) specific for either the low, neutral, or both low and neutral pH conformations of A/Memphis/1/71 (H3N2) HA^14,15^ were tested for their ability to bind to this virus after exposing it to the compounds. As conformational integrity is crucial for this test, we employed a slot blot assay in which intact virus was treated with compound dissolved in pH 7.2 saline or control solutions and then serially diluted and bound to a nitrocellulose membrane. The membranes were then probed with the MAbs and the relative intensities of the bands after development were determined (Supplementary Fig. 2a). Treatment with either MD345 or MD185 resulted in the loss of binding of the neutral pH conformation-specific MAb 241 with a concomitant gain in binding of the low pH conformation-specific 88/2 antibody, while the binding of the pH-independent MAb 244 remained unaffected (Fig. 2a, Supplementary Fig. 2b). The fact that pre-treatment of the virus with the compounds produced a binding profile mimicking pre-treatment with pH 4.9 citrate buffer confirms that the compounds are inducing the low pH conformation. It should be noted that the pH of the solutions of compounds, which was pH 7.0 after dilution in pH 7.2 saline (or pH 5.5 in unbuffered saline), was not sufficiently low in itselfself to mediate the effects we have observed, supporting the concept of creation of an acidic microenvironment at the virion surface through the binding of the compounds.

**Figure 2.**
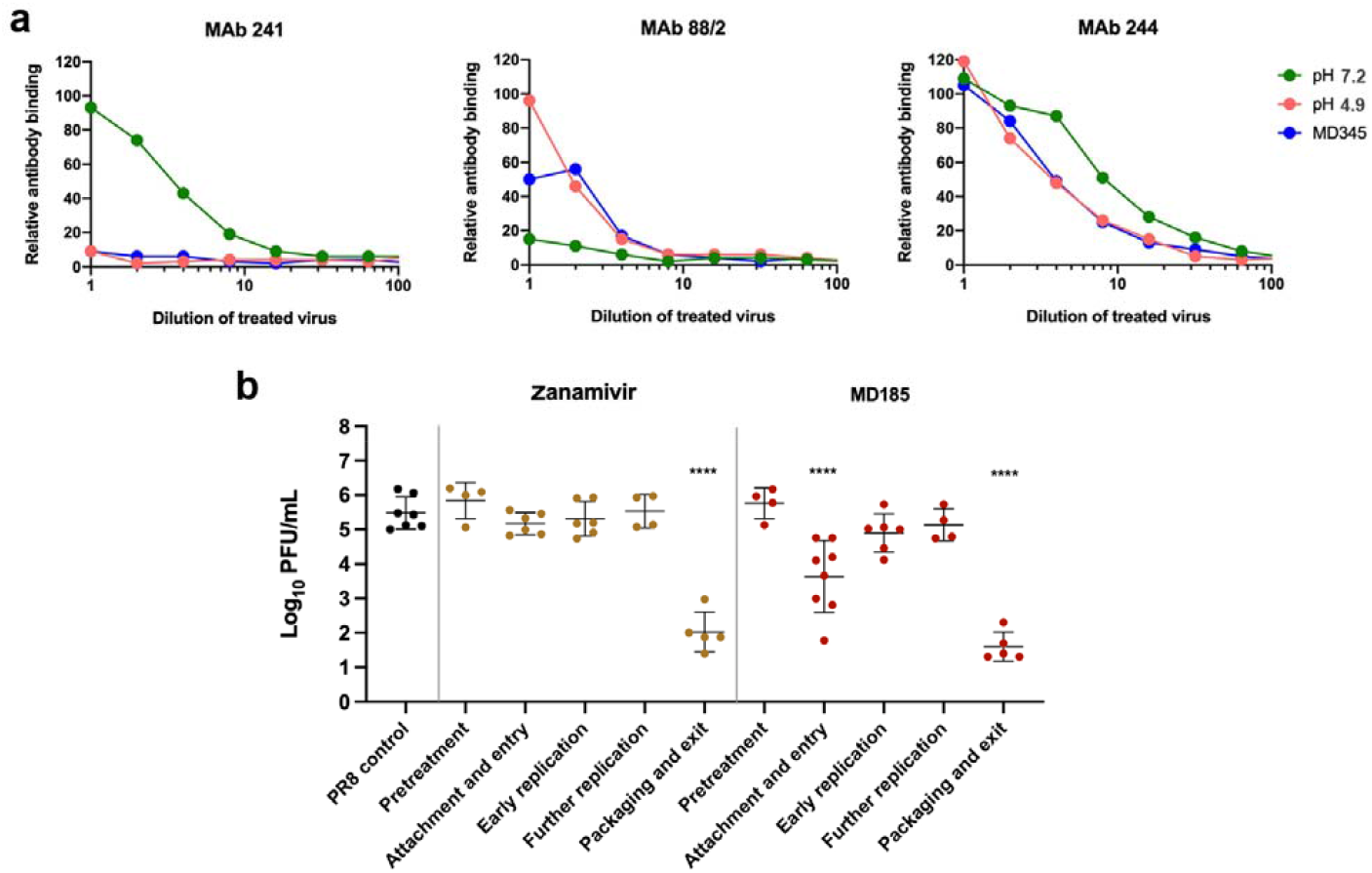
Induction of the pH-dependent conformational change in HA by the compounds and resultant inhibition of viral entry. **a**, Influenza virus was treated with MD345 (blue), pH 4.9 citrate buffer (red), or pH7.2 saline (green) and tested in a slot blot assay for binding to a panel of HA-specific MAbs. After serial dilution, the treated virus was bound to nitrocellulose and probed with: MAb 241, specific to the neutral pH conformation of HA (left panel); MAb 88/2 specific for the low pH-induced conformation (middle panel); or MAb 244, which can bind to either conformation (right panel). Data shows a representative experiment, the replicates of which are compiled and shown in Supplementary Fig. 2b. **b**, Time of addition assay. MDCK cell monolayers in a 96-well plate were infected with PR8 virus at an MOI of 0.1. In PR8 control wells, the supernatants were harvested at 12 hr post infection and the amount of infectious virus was determined by plaque assay. To establish where in the virus replication cycle the compounds were acting, 5 ug of either MD185 or zanamivir were added to additional wells at different intervals either before or after infection with PR8 virus(t=0 hr) to capture the different stages of replication indicated. At the end of the treatment interval, the cells were thoroughly washed, media was replaced and the infectious process allowed to continue to t=12 hr when supernatants were again harvested and infectious virus yields were determined by plaque assay. Data represent combined results from 5 individual experiments. ****, significantly different (P<0.0001) than the PR8 control, one-way ANOVA.

As a crude surrogate of how inducing the low pH conformation in the HA of the virus impacts its ability to interact with receptors, the compounds’ capacity to inhibit hemagglutination of chicken red blood cells was examined (Supplementary Fig. 3). In contrast to zanamivir, which failed to inhibit hemagglutination, both compounds showed marked inhibition, with MD345 being much more active. From these data, it could be predicted that the compounds modulate infection by preventing both entry, through inactivation of the HA by the effector domain, and exit, via the enzyme inhibition of the NA anchor domain. To test this we performed time-of-addition studies in MDCK cells infected with the highly virulent A/Puerto Rico/8/34 (PR8) laboratory strain over a single cycle of replication (Fig. 2b). MD185 or zanamivir was added to the cells at discrete intervals and then removed by thorough washing. In contrast to zanamivir, which had an inhibitory effect only when added at the time of viral exit from the cells, we observed a two-log reduction in viral titer when MD185 was present together with the virus during the adsorption phase, implying inhibition of viral entry. The anchor domain of the compound appeared to function analogously to zanamivir in its inhibition of virus egress.

**Figure 3.**
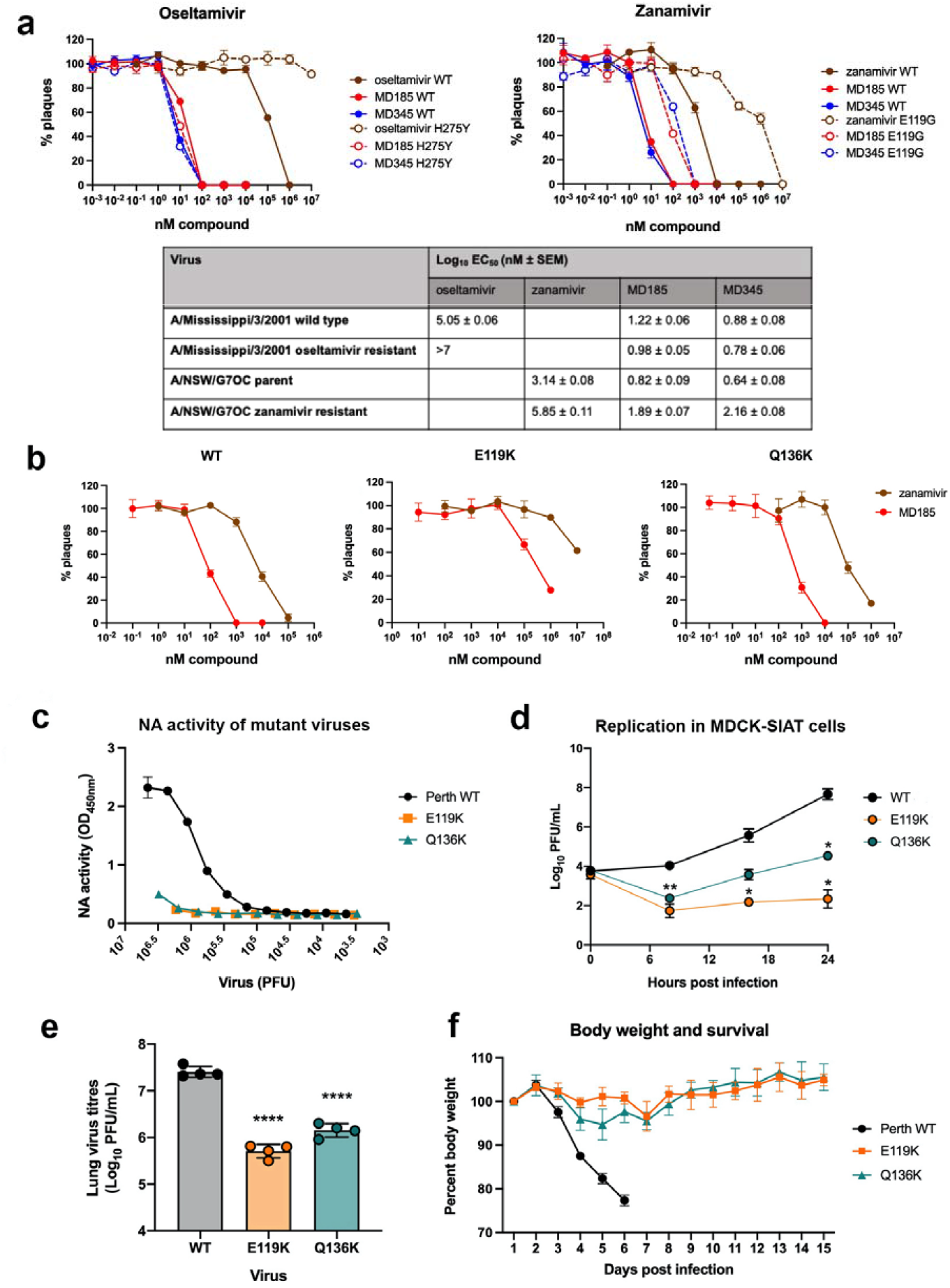
Activity of compounds against escape mutants. **a**, Activity of compounds against viruses resistant to licensed NA inhibitors. Left panel: plaque reduction assay of A/Mississippi/3/017/06/2013 WT (solid lines) and its oseltamivir-resistant mutant NA H275Y (dashed lines) by MD185 (red), MD345 (blue) or oseltamivir (brown). Right panel: plaque reduction assay of A/NWS/G70C H1N9 parent virus (solid lines) and its zanamivir-resistant mutant NA E119G (dashed lines) by MD185 (red), MD345 (blue) or zanamivir (brown). The Table insert shows the log EC_50_ of the compounds against each virus with the standard error of 3 replicate determinations. **b**, Activity of compounds against mutants with reduced susceptibility to MD185. Plaque reduction of A/Perth/265/09 WT (left panel) and its mutants NA E119K (middle panel) and Q136K (right panel) by MD185 (red) and zanamivir (brown). Data represents mean and standard error of triplicate samples of a representative of three individual experiments. The means of the log EC_50_ ± SEM for the 3 experiments for zanamivir and MD185 respectively are 3.54 ± 0.11 nM and 1.28 ± 0.37 nM for WT virus, 7.15 ± 0.13 nM and 4.92 ± 0.52 nM for NA E119K, and 5.05 ± 0.36 nM and 2.38 ± 0.31 nM for Q136K. **c**, NA activity of viruses with reduced susceptibility to MD185. The NA enzyme activity of A/Perth/265/09 WT (black), mutant E119K (orange) and mutant Q136K (teal) was assessed using fetuin as a substrate. Data shows a representative of 3 independent experiments. **d**, MDCK-SIAT1 cells were infected with 3000 PFU wild type A/Perth/265/09 virus (black), mutant E119K (orange) or mutant Q136K (teal). Supernatants were sampled at 8, 16 and 24 hr and assayed by plaque formation in MDCK cells. The mean and SEM of combined data from two individual replicate experiments each performed in duplicate are shown. Significantly different (** P<0.01, * P<0.05) from WT vírus, 2-way ANOVA. **e**, Mice (N=4) were infected with 500 PFU wild type A/Perth/265/09 virus (black), mutant E119K (orange) or mutant Q136K (teal) and the lungs sampled three days later, homogenized and assayed for infectious vírus by plaque formation in MDCK cells. Significantly different (**** P<0.0001) from WT vírus, one-way ANOVA. **f**, a second set of mice was infected as in **e** and weight loss and clinical signs monitored at least once daily. All mice infected with the wild type vírus rapidly lost weight and were euthanized on day 6 at the humane endpoint. Shown is the mean and SEM of 4 mice per group.

When evaluating any new influenza antiviral it is critical to determine its efficacy against known resistant viruses and additionally to determine whether fit, resistant mutants are readily selected after prolonged exposure to the inhibitor. Both compounds were tested for inhibition in a plaque reduction assay of the H1N1 A/Mississippi/3/01 wild type virus and the corresponding virus with an H275Y mutation in the NA that rendered it resistant to the most commonly used NA-inhibitor, oseltamivir^16^ (Fig. 3a). Against the wild type virus, MD185 was 7000-fold and MD345 was15,000-fold more potent than oseltamivir, and while the H275Y mutation conferred >>200-fold resistance to oseltamivir, neither MD185 or MD345 showed significant differences in inhibition of the mutant and wild type viruses. Since the anchor domain is a zanamivir-like molecule we also tested efficacy against the A/NWS/G70C parent virus and its E119G zanamivir-resistant mutant^17^ (Fig. 3a). In the plaque reduction assay both compounds were > 200-fold more potent than zanamivir against the parent virus, and while the E119G substitution reduced sensitivity to zanamivir by approximately 500-fold, there was only a 12-fold reduction in sensitivity to MD185 and 33-fold reduction to MD345.

To try to rapidly select for resistant variants, A/Perth/265/09 (pdmH1N1) and A/Fukui/45/04 (H3N2) viruses were passaged at a high multiplicity in the EC_90_ (10 ng/ml) of MD185. After four passages no resistant variants were isolated from the H3N2 virus, or from either virus passaged in the absence of inhibitor. However, mixed plaque phenotypes were seen in the MD185-passaged pdmH1N1 virus (Supplementary Fig. 4), and two small plaques were picked and amplified for characterization. One of these was found to possess a Q136K NA substitution, a mutation shown previously to be selected in MDCK cell culture in the absence of drug pressure ^18^, although no such change was observed in viruses passaged in the no drug/control arm of this experiment. The second had an E119K NA substitution accompanied by a G172E substitution in the HA. In a plaque reduction assay (Fig. 3b), MD185 was around 200-fold more potent than zanamivir against the wild type virus. The E119K and Q136K mutations reduced inhibition by zanamivir by over 3,000- and 30-fold respectively. Although sensitivity to MD185 was also reduced, it was still greater than 150-fold more effective than zanamivir against both mutants. Thus even with viruses containing substitutions conferring reduced NA inhibitor sensitivity, the acidic effector domain of the compounds provided robust anti-viral activity.

**Figure 4.**
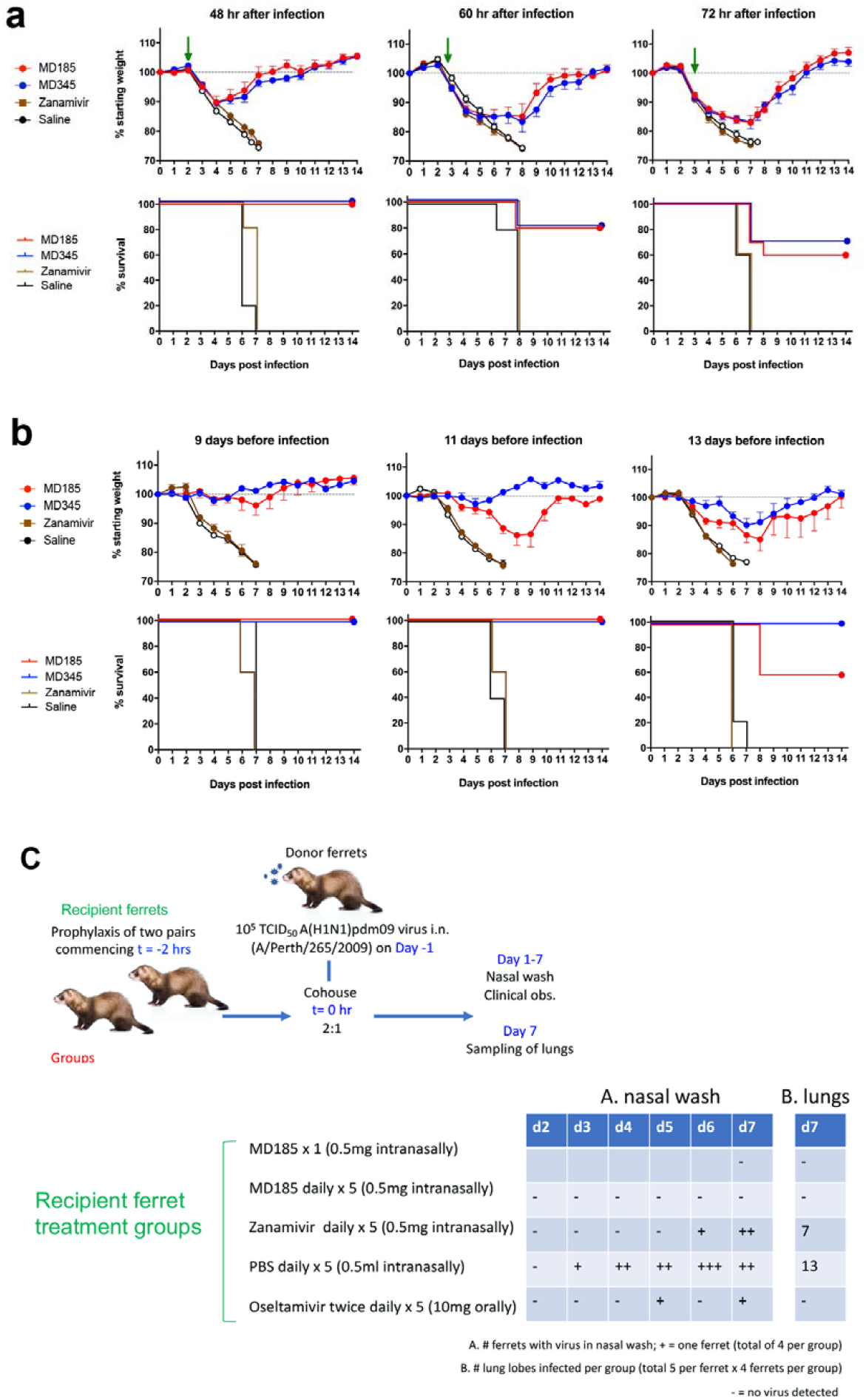
*In vivo* activity of the compounds. **a**, Ability of the compounds to reduce viral loads in infected mice. Mice were infected with a lethal dose (500 PFU) of PR8 virus then treated intranasally with 40 μg MD185 (red), MD345 (blue), zanamivir (brown) or saline (open black). Upper panels show mean and SEM of weights of the mice in each group measured daily for 14 days after infection, the individual data for which is shown in Supplementary Fig. 5. Data are expressed as a percentage of the starting weights. Lower panels plot the percent survival of the same mice when euthanized at the pre-determined humane endpoint. Green arrows indicate time of treatment at 48, 60 or 72 hr after infection. **b**, Ability of the compounds to prevent infection-induced weight loss and death in mice. Mice were given 5 μg MD185 (red), MD345 (blue), zanamivir (brown) or saline (open black) intranasally either 9,11 or 13 days before infection with a lethal dose (500 PFU) of PR8 virus. Data are presented as in **a**. Individual mouse data for this experiment are shown in Supplementary Fig. 6. **c**, Ability of compounds to impact viral growth in the respiratory tract of ferrets. Donor ferrets were infected and then cohoused with treated recipient ferrets as outlined. Nasal washings from the recipient ferrets were taken daily after exposure to the donor ferrets, except for the group treated with a single dose of MD185 that had nasal washings on day 7 only. Individual lung lobes of recipient ferrets were sampled on day 7. These and the nasal wash samples were assayed for infectious vírus by a CPE assay in MDCK cells. For nasal washings (A), + represents a vírus positive ferret, - indicates no vírus detected, no symbol is not undergoing nasal washing. For the lungs (B), the data shown is the number of infected lobes out of a total of 20/group. - indicates no vírus detected. Titres for individual ferrets are shown in Supplementary Fig. 7.

Further characterization demonstrated that the NA activity of the mutants was barely detectable (Fig. 3c), suggesting that the normal functions of NA in penetration through the sialylated mucosal layer and efficient exit from cells would be compromised, reducing replication^19^. To test this, we infected MDCK-SIAT 1 cells, which over express sialyl-α2,6-galactose moieties on the cell surface, and assessed virus yields over a 24 hr period (Fig 3d). Both mutants showed decreased production of virus compared to the A/Perth/265/09 WT virus when replicating in the presence of a high density of cell surface receptor. At 24 hr after infection of the cells, approximately 3- and 5-log_10_ reductions in virus yields were observed for the Q136K and E119K mutants respectively. These data suggest that replication may also be compromised *in vivo* as seen with other mutants with low NA activity^20^. This was confirmed by the reduced pulmonary virus titers observed in mutant virus-compared to wild type virus-infected mice (Fig. 3e). Likewise, only mild weight loss (Fig. 3f) and complete survival was observed in mice infected with a dose of mutant virus, the equivalent of which was lethal using the wild type parent strain.

Finally, we tested the *in vivo* activity of the compounds. Mice were infected with a lethal dose of PR8 virus followed by intranasal treatment with a single 40 μg (2 mg/kg) dose of either compound or zanamivir at 48, 60, or 72 hr post infection (hpi) (Fig. 4a, Supplementary Fig. 5). When treatment was initiated within 48 hpi, both MD185 and MD345 caused a significant reduction in morbidity and completely eliminated the need for euthanasia at the humane endpoint. Only when treatment was delayed to 60 hpi was a reduction in protection observed, with an 80% survival rate with both compounds. Even at 72 hpi, at a time point where the mice are already displaying clinical symptoms and significant weight loss, MD185 and MD345 were able to rescue 60 and 70% respectively of the animals treated. In humans, successful therapy at such delayed time points is an important factor, given that many patients do not present until more than 48 hr after development of symptoms when currently licensed antivirals may prove ineffective^21^.

To test the prophylactic potential of the compounds, mice were given a single intranasal 5 μg (0.25 mg/kg) dose of either MD185, MD345 or zanamivir at 9, 11, or 13 days before infection with a lethal dose of PR8 (Fig. 4b, Supplementary Fig. 6). Mice survived when given MD185 or MD375 up to 11 days before infection. Although the MD185-treated group began to show weight loss when the time interval between administration of compound and infection was 11 days and mortality when the interval was increased to 13 days, with MD345 no clinical signs were observed with a 13-day prophylactic interval and even then all mice were protected from death. Notably, using only one fortieth of the dose, MD345 extended the duration of the protective effect beyond that observed with the most potent of zanamivir dimers ^22^. Overall, these data would indicate that while the *in vivo* therapeutic and *in vitro* efficacy are similar between the two compounds there appears to be a greater prophylactic capacity with MD345 compared to MD185.

While the compounds were effective at preventing severe disease in mice, the murine model, which involves deposition of virus throughout the respiratory tract, does not accurately represent how influenza is acquired in nature. To explore the effectiveness of the compounds in a more representative model, outbred ferrets were given either 0.5 mg MD185 intranasally either once only or once per day for five days, 0.5 mg zanamivir intranasally once daily for five days, 5 mg oseltamivir orally twice daily for five days or PBS as a placebo, starting two hr before co-housing with A/Perth/265/09 (pdmH1N1) virus-infected donors (Fig. 4c). All donor ferrets were confirmed to be shedding from day one post infection (Supplementary Fig. 7a). Of the ferrets cohoused with the infected donors, 3 of 4 in the placebo group had virus in the daily nasal washings detectable initially between 3-6 days after cohousing and all had virus in at least one lung lobe when sampled on day 7 post cohousing (Fig. 4c, Supplementary Fig. 7b-d). Two of four zanamivir-treated ferrets had virus in both nasal washings and lung lobes while a third tested positive only in the lungs. Of the three ferrets treated continuously with oseltamivir, one had virus in the day 5 and day 7 nasal washings but none had virus in the lungs. In contrast, treatment with either a single dose or 5 once-daily doses of MD185 completely prevented the establishment of infection in both the upper and lower respiratory tract. Notably, the single treatment with MD185 achieved greater protection than oseltamivir, even when the latter in a single day is given at a dose twenty times greater by weight and 119 times greater by molarity.

These *in vivo* data highlight the effectiveness of the compounds after a single small dose, even for long-duration prophylaxis as with MD345, indicating a potential role for post-exposure prophylaxis or for prevention in a pandemic context. Single dose oral treatment of influenza infection is an advantage of the most recently approved influenza antiviral drug, baloxavir marboxil, a viral endonuclease inhibitor. However, the drug is prone to selection of resistant mutants^23^ and these may be sufficiently fit to spread amongst individuals ^24^, emphasising the need for continued exploration of new candidates.

By targeting the acidic effector domain to the virus surface we were able to achieve prolonged and efficient inactivation of virus with a mode of action consistent with the creation of an acidic microenvironment at the virion surface. Although an environmental approach in the treatment of influenza has been attempted before by Rennie *et al*.^25^ who showed that a low pH nasal spray was effective at treating influenza virus-infected ferrets, this therapy required a high concentration and had a very limited residence. The highly successful application of our strategy to influenza virus should provide the impetus to use this as a blueprint for the construction of a new class of antiviral compounds against the spectrum of viruses, including *Corona-, Rhabdo*-, *Arena*-, *Bunya*-, *Flavi*-, *Henipa*-, *Hepadna*-, *Pox*- and *Togaviridae* families, that use low pH as a trigger for endosomal membrane fusion^26^ as well as other pathogens for which inhospitable microenvironments and appropriate anchors can be envisaged.

## Methods

### Synthesis of MD185 and MD345

The synthesis of the compounds is shown in the Supplementary Methods

### Viruses and cells

Madin-Darby canine kidney (MDCK) cells were grown in Roswell Park Memorial Institute (RPMI)-1640 medium (Sigma Aldrich, Castle Hill, Australia) supplemented with 10% heat-inactivated fetal calf serum (FCS, Life Technologies, Mulgrave, Australia), 2 mM L-glutamine (Sigma-Aldrich), 2 mM sodium pyruvate, 24 μg/ml gentamicin (Pfizer, West Ryde, NSW, Australia), 50 μg/ml streptomycin and 50 IU/ml penicillin unless otherwise stated. MDCK-SIAT 1 cells were obtained from the WHO Influenza Centre for Reference and Research, Melbourne and were grown in the same medium. The influenza viruses used in this study were obtained from the collection at the Peter Doherty Institute. Stocks of A/Puerto Rico/8/34 (H1N1), A/Memphis/1/71 (H3N2), A/California/7/09 (pdmH1N1), A/Switzerland/9715293/2013 (H3N2) and B/Malaysia/2008 were amplified in 10-day embryonated hen’s eggs and stocks of A/Perth/265/09 (pdmH1N1), A/Fukui/45/04 (H3N2), A/ Mississippi/3/01 (H1N1) and its H274Y oseltamivir-resistant mutant and A/NWS/G7OC (H1N9) and its E119G zanamivir-resistant mutant were amplified in MDCK cells.

### Assay of infectious virus by plaque formation

Viruses were enumerated by plaque assay performed in duplicate wells in TC-6 plates (Corning) of confluent MDCK cell monolayers overlaid with 3 ml of Leibovitz’s L-15 medium with L-glutamine (Gibco), 0.2 mM HEPES buffer, 0.9% (w/v) Type 1 agarose (Sigma) and 1 μg/ml L-1-tosylamido-2-phenylethyl chloromethyl ketone (TPCK)-treated trypsin (Worthington Biochemicals). Plaque forming units (PFU) were counted by eye after 3 days.

### Viral cytopathic effect (CPE)-reduction assay

This assay was used for the initial screening of compounds against a range of virus strains. MDCK cells were maintained in Minimal Essential Medium (Gibco, Thermo Fisher) supplemented with 1 mM non-essential amino acids, 50 U/ml penicillin, 50 μg/ml streptomycin, 1 mM sodium pyruvate and 10% fetal bovine serum. For assay of compounds, MDCK cells were grown to confluency in 96-well microtiter trays. Media was removed and 46 μl of virus at 10-fold above the 50% tissue culture infectious dose (TCID_50_) was added to each well. Plates were incubated at 37°C for 1.5 hr after which the viral inoculum was removed and replaced with medium containing trypsin (TPCK-treated). Test compounds were diluted in 1:3 series in the wells. Control wells contained medium in place of compound. After 48-96 hr, the medium was removed and replaced with medium without trypsin. Viable cells were detected after the addition of CellTiter 96 Aqueous One Solution reagent (Promega) by measurement of absorbance at 490 nm, according to the manufacturer’s instructions.

### Plaque reduction assays

MD185 and MD345 were dissolved in RPMI at a concentration of 200 nM and zanamivir was reconstituted to 2000 nM before each sample was 10-fold serially diluted to 0.001 nM. Influenza virus at 1×10^3^ PFU/ml in RPMI was mixed 1:1 with compound dilutions and preincubated at room temperature for 1 hr. Confluent MDCK monolayers in six-well plates were washed with RPMI and 200 μl per well of the virus/compound dilutions were added to triplicate wells. The plates were left at 37°C for 1 hr before being overlaid with medium containing agarose as above. Three days post infection, readily visible plaques were counted. Notably, as the expected mode of action of the compounds is to inhibit virus entry, compounds were pre-incubated with virus rather than being added to the overlay medium as would be done to assess inhibitors of the NA, where a reduction in plaque size rather than number is typically measured.

### Neuraminidase assay with fetuin substrate

A 96-well Maxisorp plate (Thermo Fisher, Denmark) was coated with 100 μl of 125 μg/ml fetuin (Sigma Aldrich) in phosphate buffered saline (PBS) and left overnight at 4°C. Virus was serially diluted to 5×10^7^ PFU/ml in stability buffer (0.9% NaCl, 1% bovine serum albumen, 10 mM CaCl_2_ and 5 mM MgCl_2_) containing 0.5% Triton X-100 to separate the activity of the NA from the binding capacity of the HA, and left at room temperature for 20 min. The fetuin plate was washed three times with PBS and 100 μl/well of each dilution was added in duplicate before the plate was incubated at 37°C for 16 hr. The plate was washed six times in PBS with 0.05% Tween-20 (PBST) then incubated with horse radish peroxidase-conjugated peanut Arachis Hypogaea lectin (Cosmo Bio) for 1 hr at room temperature. Further washes in PBST were performed before 100 μl/well of SureBlue TMB substrate (Seracare) was added and left at room temperature for 20 min. The reaction was stopped with 50 μl/well of 1 M HCl and read at 450 nm.

For NA inhibition assays, MD185, MD345, or zanamivir (45 μM) were 10-fold serially diluted in stability buffer then incubated 1:1 with virus (5×10^7^ PFU/ml) for 20 min at room temperature prior to adding to the washed fetuin plate.

### Neuraminidase inhibition assay with MUNANA substrate

NA activity was measured using the MUNANA fluorescence-based assay as described previously^27^. The assay measures the increase in fluorescence of free methylumbelliferone released from MUNANA after cleavage by NA. Viruses, standardized to 100 fluorescence units, were preincubated in microtiter plates for 30 min at 37°C with serial 2-fold dilutions of compound. MUNANA (Carbosynth) was then added, the assay mixtures were incubated for 1 hr at 37°C, and the assays were stopped by the addition of 0.2 M Na_2_CO_3_. Fluorescence was measured in a BMG FLUOstar Optima reader with an excitation wavelength of 365 nm and an emission wavelength of 450 nm.

### Slot Blot assay

A/Mem/l/71-Bel/42 (H3N1) (3.4×10^7^ PFU/ml) was diluted 1 in 10 in either pH 7.2 saline, pH 4.9 10 mM sodium citrate buffer, or 340 μM of MD345 or MD185 in pH 7.2 saline and left at room temperature for 2 hr. Each virus sample was then 2-fold serially diluted in saline before 50 μl was added to triplicate wells of a Minifold 2 slot blot apparatus^28^. After binding, filters were removed, cut into 3 replicate strips and blocked with 4 ml 1% casein in PBS for 10 min. Separate strips were then reacted for 1 hr with one of three monoclonal antibodies: MAb 88-2 which detects only the low pH conformation of HA, MAb 241 which detects only the neutral pH conformation, or MAb 244 which can bind to both conformations. After washing, an HRP-conjugated rabbit anti-mouse antibody (Dako) was applied for 1 hr, strips were washed and SureBlue TMB substrate (Seracare) with 1% dextran sulphate was added to each strip. After developing, the strips were rinsed with water, dried and scanned with a docuprint CM415 AP Fuji xerox scanner. Band density was analysed using ImageJ software.

### Hemagglutination inhibition

PR8 virus (2.6 × 10^4^ PFU) was preincubated in Eppendorf tubes with either saline or the indicated amounts of MD185, MD345 or zanamivir. After 2 hr, serial 2-fold dilutions of the mixtures were performed in round-bottomed 96-well microtiter plates and an equivalent volume of 1% (vol/vol) chicken erythrocytes added. The pattern of hemagglutination was determined after 30 mins.

### Time of addition assay

Monolayers of MDCK cells in 96-well plates were infected with an MOI of 0.1 PFU/cell starting at t= -1 hr. Duplicate wells were treated with 5 μg of either MD185 or zanamivir in 100 μl RPMI-1640 media. Each treatment occurred for a period two hours representing discrete stages in the influenza virus life cycle: pre-treatment, from -4 hr to -2 hr; attachment and entry, a one-hour preincubation of virus and compound from -2hr to -1hr followed by addition to cell monolayer from -1 hr to 0 hr; early replication, from 0 h to 2 hr; late replication, from 2 hr to 4 hr; packaging and exit, from 4 hr to 6 hr post infection. Control wells were left untreated. Following treatment, cells were washed extensively to remove any trace of compound, this involved two initial washes followed by two 30 min washes in 200 μl RPMI. At 12 hpi, supernatants were collected, serially diluted and virus assayed by plaque formation in MDCK monolayers.

### Selection and sequencing of resistant mutants

Ten-fold serial dilutions of A/Perth/265/09 (pdmH1N1) and A/Fukui/45/04, (H3N2) viruses were passaged at a multiplicity of >1.0 in MDCK cells in 24-well dishes for 4 passages. MD185 was added at 10 ng/ml, and control wells included virus without any inhibitor to detect any adaptive mutations. Virus titer was determined after each passage by plaque assay, and plaque morphology was examined for phenotypic changes indicative of potential mutations, which were seen by passage 4. Potential mutants were plaque purified in MDCK monolayers and picked into 200 μl RPMI-1640 medium. Viral amplification was performed in chicken egg embryos for 48 hr at 35°C or MDCK cells for 72 hr at 37°C. RNA was extracted from viral supernatants using the RNeasy mini kit (Qiagen, Germany) followed by cDNA conversion using the Omniscript RT kit (Qiagen, Germany) according to kit instructions. DNA was amplified using a Phusion Flash PCR master mix (Thermo Fisher) with HA and NA specific primers (primers available on request). The samples were gel purified using the QIAquick gel extraction kit (Qiagen, Germany) and sequencing was performed using the BigDye Terminator v3.1 Cyle Sequencing Kit (Applied Biosystems, Massachusetts, USA).

### Mouse infection experiments

Experiments were conducted under approval from the University of Melbourne’s Animal Ethics Committee (Projects 1112112.3, 1212572.4, 1212665.4 and 1413321.3 and 1714217.2) in accordance with the Australian code for the care and use of animals for scientific purposes. Seven to eight-week-old female BALB/c were obtained from the Animal Resources Centre (ARC, Perth, Australia) and housed at the Bioresources facility at the Peter Doherty Institute. The mice were anesthetized using isoflurane and treated intranasally with 5 μg/mouse for prophylaxis or 40 μg/mouse for therapy of MD185, MD345 or zanamivir in 50 μl of 0.9% saline. At the specified times pre-or post-treatment, the animals were anesthetized again and infected intranasally with a lethal dose (500 PFU) of PR8 in 50 μl 0.9% saline. All infected animals were weighed and monitored for clinical signs at least once daily and euthanized at the predefined humane endpoint.

In some experiments, C57BL/6 mice (Peter Doherty Institute) were infected with variants of A/Perth/265/09 virus selected in the presence of MD185 or with the WT parent virus. Lungs were harvested from the mice on day 3 after infection and assayed for infectious virus by plaque formation in MDCK cells.

### Ferret infection experiments

Experiments were conducted under approval from the University of Melbourne’s Animal Ethics Committee (Project 1313040) in accordance with the Australian code for the care and use of animals for scientific purposes. Six to twelve-month-old outbred ferrets (12 female, 17 male *Mustela putorius furo*; Animalactic Animals and Animal Products, Australia) were housed at the Bioresources Facility within the Peter Doherty Institute. The day before co-housing, ten female donor ferrets were anesthetized using an intramuscular Xylazine:Ketamine injection before being infected intranasally with 10^5^ TCID_50_ A/Perth/265/2009 virus. The remaining animals were randomly allocated into one of five treatment groups: 0.5mg of MD185 given intranasally once only; 0.5 mg of MD185 given intranasally once a day for five days; 0.5 mg of zanamivir given intranasally once a day for five days; 500 μl of PBS placebo given once a day for five days; or 5mg of oseltamivir (Sequoia Research Products) given orally twice a day for five days. All treatments were performed under Ketamine:Midazolam:Medetomidine anesthesia except for the oseltamivir group where drug was administered to conscious ferrets. Two hours after the initial treatment one donor ferret was co-housed with two recipient ferrets for a 48 hr period before the donor was removed. The recipient animals were monitored daily for changes in temperature, activity or bodyweight compared to a pre-experimental baseline, while nasal washes were taken starting 24 hr post infection to determine viral titer. A nasal wash was performed on the MD185 single administration group only once on day 7 due to fears that residual compound would be washed away. Seven days after co-housing, all animals were euthanized and individual lung lobes dissected for analysis of virus titers using a CPE assay. Titers of virus in nasal washes and lung homogenates were expressed as TCID_50_/ml determined in MDCK cells.

### Statistical analysis

All statistical analyses were performed with GraphPad Prism version 9.4.1 using one-way analysis of variance (ANOVA) or, for the neuraminidase inhibition titration, two-way ANOVA, both with Sidak’s multiple-comparison tests.

