## Supplementary Table for "A new concept in antiviral drug design yields a potent influenza inhibitor"

**Supplementary Table 1 CPE reduction screening assay**

| Influenza virus |  | EC_50_ (nM) |  |  |
| --- | --- | --- | --- | --- |
| Type/Subtype | **Strain** | **MD185** (MW1852) | **MD345** (MW2201) | **Zanamivir (**MW332) |
| A/H1N1 | A/Sydney/250/1999 | 0.29 | 0.052 | >198 |
|  | A/Solomon Islands/3/2006 | 0.29 | 0.043 | >44 |
|  | A/Mississippi/03/2001 | 0.89 | 0.24 | >66 |
| A/H1N1pdm09 | A/Perth/261/2009 | 0.49 | 0.32 | 6.6 |
|  | A/California/7/2009 | 3.2 |  |  |
|  | A/Townsville/74/2011 | 0.74 | 0.041 | >15 |
| A/H3N2 | A/Sydney/5/1997 | 2.7 | 0.83 | >24 |
|  | A/Victoria/503/2006 | 6.5 | 2.2 | >198 |
|  | A/Perth/16/2009 | <0.07 |  |  |
|  | A/Victoria/170/2012 | 2.7 | 0.53 | >596 |
| A/H5N1 | A/Duck/Minnesota/1525/1981 | <0.07 |  |  |
|  | A/Hong Kong/213/2003 | <0.17 |  |  |
| A/H7N9 | A/Anhui/1/2003 | <0.17 |  |  |
| B | B/Florida/4/2006 | <0.07 |  |  |
|  | B/Townsville/2/2011 | 1.7 | 0.24 | >1789 |
