## Supplementary Methods for "A new concept in antiviral drug design yields a potent influenza inhibitor"

### Abbreviations

The following abbreviations are used in the description of the synthesis of compounds: methyl (Me), ethyl (Et), n-propyl (nPr), iso- propyl (iPr), n-butyl (nBu), tert-butyl (tBu), n-hexyl (nHex), cyclohexyl (cHex), phenyl (Ph), methoxy (MeO), ethoxy (EtO), trimethylsilyl (TMS), tert-butyloxycarbonyl (Boc), and acetyl (Ac), methanol (MeOH), ethanol (EtOH), diethyl ether (Et20), ethyl acetate (EtOAc), triethylamine (TEA), dichloromethane (methylene chloride, DCM, $CH_2C1_2$), trifluoroacetic acid (TFA), trifluoroethanol (TFE), dimethylformamide (DMF), sodium sulphate ($Na_2S0_4$), tetrahydrofuran (THF), meta-chloroperoxybenzoic acid (mCPBA), hexamethyldisilazane sodium salt (NaHMDS), 0-(7-azabenzotriazol-1-yl)-N,N,N',N'-tetramethyluronium hexalluorophosphate (HATU), O-(benzotriazol- 1-y l)-N,N,N',N'-bis(tetramethylene)uronium hexalluorophosphate (HBTU), dimethylsulfoxide (DMSO), magnesium sulphate ($MgS0_4$), sodium hydrogen carbonate ($NaHC0_3$), tert-butanol (i-BuOH), l-ethyl-3-(3-dimethylaminopropyl) carbodiimide hydrochloride salt (EDC1.HCl), tetra-n-butylammonium fluoride (TBAF), N,N-diisopropylethylamine (DIPEA), tert-butyldimethylsilyl (TBDMS), 1- hydroxybenzotriazole (HOBt), trans-dichlorobis(triphenylphosphine)palladium(II) ($PdCl_2(PPh_3)_2$), tetrakis(triphenylphosphine)palladium(0) ($Pd(PPh_3)_4$) tris(dibenzylideneacetone) dipalladium(O) ($Pd_2(dba)_3$), tri-t-butyl phosphonium tetrafluoroborate ($i\text{-}Bu_3PH.BF_4$), 4,5-bis(diphenylphosphino)-9,9-dimethylxanthene (Xantphos), triphenylphosphine ($PPI1_3$), diisopropyl azodicarboxylate (DIAD), pyridinium chlorochromate (PCC), borane dimethylsulfide (BMS), titanium isopropoxide ($TiOiPr_4$), sodium triacetoxyborohydride ($NaBH(OAc)_3$), sodium cyanoborohydride ($NaBH_3(CN)$), sodium borohydride ($NaBH_4$), ammonium chloride ($NH_4C1$), chloroform ($CHCl_3$), manganese dioxide ($Mn0_2$), potassium carbonate ($K_2C0_3$), 1,2-dichloroethane (DCE), sodium azide ($NaN_3$), sodium nitrite ($NaN0_2$) di-tert-butyl dicarbonate ($Boc_20$), and S-Acetamidomethyl (Acm).

### Synthesis of anchor compound Zn’

The synthesis of Zn’ first required the production of N-Boc-1,9-diaminononane. 1,9-Diaminononane (1 g) was dissolved in a mixture of 50 ml ethanol and 50 ml water, then di-i-butyldicarbonate (1.38 g) was added at room temperature (rt). The whole mixture was stirred at rt for 16 hrs to afford a white suspension. This suspension was filtered off. The solid was di-Boc-l,9-diaminononane (488 mg) after air drying. The filtrate was extracted with dichloromethane. The organic extracts were combined and washed with 50 ml of water, then stirred with 10 g anhydrous $Na_2S0_4$ at rt overnight. The organic suspension was filtered. The filtrate was vacuum evaporated to dryness to afford N-Boc-l,9-diaminononane 809 mg (49.5% yield [yd.]) as a colourless solid. MS 295(M+1).

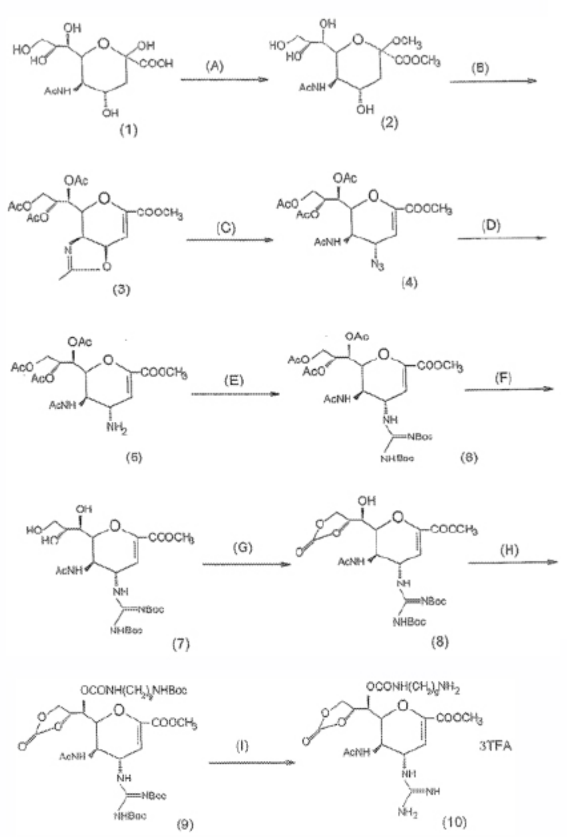

The synthesis of Zn' then proceeded according to the steps (A-I) shown in the figure above involving compounds 1-10.

Step A: Sialic acid (compound 1) (5 g, 16.18 mmoles) and Dowex 50 x 8($H^+$) resin (10 g) were stirred in anhydrous methanol (400 ml) at 60-62 °C for 48 hrs. The mixture was filtered off. The filtrate was vacuum evaporated to dryness to afford compound (2) as a white solid 4.5 g (13.25 mmoles, 82.5% yd.). MS 338(M+1).

Step B: Compound (2) (4 g, 11.87 mmoles) was stirred with acetic anhydride (40 ml, d 1.08, MW 102.29, 423 mmoles) and sulphuric acid (2 ml). The mixture was then stirred at 32-35°C for 72 hr in an oil-bath. The reaction mixture was added dropwise to a stirring mixture of $Na_2C0_3$ (51 g) in water (280 ml) and ethyl acetate (14 ml) at an ice- bath. Afterwards, the mixture was stirred for an additional 1.5 hr in an ice-bath, and subsequently extracted with ethyl acetate (400 ml, 270 ml). The ethyl acetate extracts were combined and washed with 10% $NaHC0_3$ solution (270 ml x 2), saturated NaCl solution (270 ml x 2) and dried over anhydrous $Na_2S0_4$ overnight and filtered off. The filtrate was vacuum evaporated to dryness to afford compound (3) 4.78 g (11.57 mmoles, 97.5% yd.). MS 414(M+1). The oily residue turned into an off white solid after storage in a desiccator over $P_2Os$ for 3 days.

Step C: Compound (3) (3.47 g, 8.4 mmoles) was dissolved in tert-BuOH (25 ml). To this solution was added azidotrimethylsilane (1.81 ml, 13.63 mmoles). The whole mixture was stirred at 80-82°C under argon for 24 hrs. The reaction mixture was diluted with ethyl acetate (100 ml), then stirred with 0.9 g $NaN0_2$ in 25 ml of water and adjusted to pH 2 with 5N HCl over a period of 1 hr at rt. The two phase mixture was separated and the aqueous layer extracted with ethyl acetate (50 ml). The organic extracts were combined and washed successively with water (25 ml x 2), 6% $NaHC0_3$ solution (25 ml), water (25 ml x 3), then dried over $Na_2S0_4$. The filtrate was vacuum evaporated to dryness to afford compound (4) 3.17 g (6.95 mmoles, 82.7% yd.) as an oily substance. MS 457(M+1), 479(M+Na), 925(2M+Na).

Step D: Compound (4) (3.15 g, 6.91 mmoles) was dissolved in methanol (100 ml) and toluene (70 ml) . The solution was placed under vacuum to remove air (oxygen), then backfilled with argon. To this mixture was added Pd/C (10%) (616 mg), then placed under vacuum to evacuate argon which was subsequently replaced with hydrogen ($H_2$). The hydrogenation was carried out at rt for 2 hr and the catalyst was subsequently filtered off. The filtrate was vacuum evaporated to dryness to afford compound (5) 2.82 g (6.55 mmoles, 94.7% yd.) as an off white solid. MS 431(M+1).

Step E: Compound (5) (2.80 g, 6.51 mmoles) was dissolved in anhydrous acetonitrile (15 ml). To this solution was added N,N'-Di-Boc-1H-pyrazole-1-carboxamidine [Bis(Boc)PCH] (3.03 g, 9.76 mmoles). The whole mixture was stirred under argon at rt for 40 hr. The reaction mixture was vacuum evaporated to dryness. The residue was dissolved in ethyl acetate (4 ml), then diluted with hexane (4 ml) and subjected to flash column chromatography, firstly washed with hexane (150 ml), then with solvent (ethyl acetate: hexane 1: 1) (150 ml) and finally eluted with solvent (ethyl acetate:hexane 1: 1) Vacuum evaporation of the collected fractions gave the product compound (6), 3.23 g (4.8 mmoles, 73.7% yd.) as a white solid. MS 673(M+1).

Step F: Compound (6) (2.8715 g, 4.273 mmoles) was dissolved in anhydrous methanol (22 ml). To this solution was added $NaOCH_3$ (4.9134 mg, 0.2136 mmoles) under argon. The whole mixture was stirred under argon at rt for 2.5 hr, then adjusted to pH 6.5 with Dowex50 x 8($H^+$) resin, which was subsequently filtered off. The filtrate was vacuum evaporated to dryness to give compound (7) 2.2024 g (4.0337 mmoles, 94.4% yd.) as a white solid. MS 547(M+1).

Step G: Compound (7) (2 g, 3.66 mmoles) was dissolved in anhydrous acetonitrile. To this solution was added 1,1'-carbonyldiimidazole (714 mg, 4.403 mmoles). The whole mixture was stirred under argon at 20-30°C for 18 hr. It was vacuum evaporated to dryness, then the residue was subjected to flash column chromatography, firstly washed with hexane (150 ml), then developed with solvent (ethyl acetate:hexane 2: 1). The fractions at $R_f$ value of 0.5 (ethyl acetate:hexane 2: 1) were combined and vacuum evaporated to dryness to afford compound (8) 1.35 g (2.36 mmoles, 64.5% yd.) as a white solid. MS 573(M+1).

Step H: Compound (8) (1.3 g, 2.27 mmoles) was dissolved in anhydrous pyridine (12.5 ml). To this solution were added 4-nitrophenyl chloroformate (503.3 mg, 2.497 mmoles) and 4-dimethylamino-pyridine (808.5 mg, 6.62 mmoles). The whole mixture was stirred at 30°C under argon for 7 hr. To this reaction mixture a solution of N-Boc-1,9-diaminononane (690 mg, 2.67 mmoles) and 4-dimethylamino pyridine (166 mg, 1.36 mmoles) in anhydrous pyridine (4 ml) was added. The reaction mixture was stirred under argon at 30°C for 16 hr, then vacuum evaporated to remove pyridine. The residue was partitioned between ethyl acetate (300 ml) and water (50 ml) containing 2.45 ml of 5N HCl, washed successively with water (50 ml x 2), 2% $NaHC0_3$ solution (50 ml x 6), water (50 ml x 2), dried over anhydrous $Na_2S0_4$ and filtered. The filtrate was vacuum evaporated to dryness to give 2.25 g of an oily substance. It was subjected to flash column chromatography (ethyl acetate:hexane 1.5: 1 as eluent). The fractions (at a Rf value of 0.27 TLC, ethyl acetate:hexane 1.5: 1 as developing solvent) were combined and vacuum evaporated to dryness to afford compound (9) 1.057 g (1.234 mmoles, 54.4% yd.) as a white solid. MS 857(M+1), 879(M+Na). 1H-NMR (DMSO-$d_6$) 5(ppm) 11.42 (1H, s, guanidine NHBoc), 8.25 (1H, d, NH-4), 7.95 (1H, d, AcNH), 7.16 (1H, t, OCONH), 6.75 (1H, t, nonyl NHBoc), 5.80

(1H, s, H-3), 4.92 (2H, m,H-7, H-8), 4.75 (1H, m, H-4), 4.31- 4.62 (4H, m, H-5, H-6, H-9, H-9'), 3.74 (3H, s, $COOCH_3$), 2.90 (4H, m, $NHCH_2(CH_2)_7CH_2NH$), 1.86 (3H, s, $NHCOCH_3$), 1.49 (9H, s, Boc), 1.38 (18H, s, Boc), 1.2-1.6 (14H, m, $NHCH_2(CH_2)_7CH_2NH$).

Step I: Compound (9) (1 g, 1.168 mmoles) was dissolved in a mixture of trifluoro acetic acid (TFA) (36 ml) and methylphenylether (Anisole) (3.9 ml) in dichloromethane ($CH_2C1_2$) (36 ml). The whole mixture was stirred at 25 °C for 2 hr and 40 min, then it was vacuum evaporated at 35°C for 2 hrs. The residue was stirred in hexane (100 ml) at rt overnight, the hexane was decanted and fresh hexane (60 ml) was added, and stirring was continued for 4 hrs at rt. The hexane was then removed. The residue was dissolved in $CH_2C1_2$ (10 ml) and evaporated to dryness at 35-40°C. The residue was dissolved in water (25 ml). The aqueous solution was freeze dried to afford compound (10) 1.026 g (1.143 mmoles, 97.8% yd.) as a white foam of $TFA_3Z_n$' salt. MS 557 (M+I) [MW of $Z_n$' = 556, $TFA_3Z_n$' = 898].

**Synthesis of Anchor-Backbone compound PYR(Zn')2**

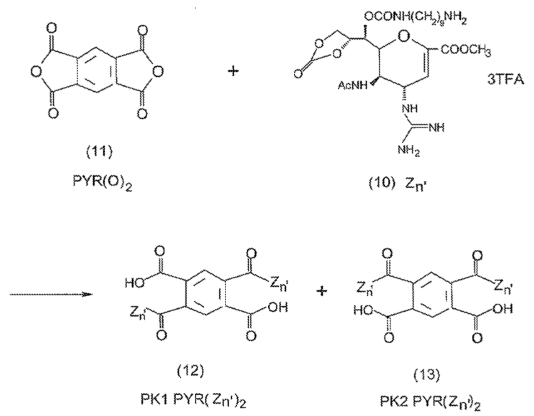


Synthesis of PYR(Zn')2 proceeded according to the step shown in the figure above involving compounds 11-13.

To a solution of pyromellitic dianhydride (compound 11) (27.25 mg, 0.125 mmoles) in anhydrous DMF (2 ml) $Z_n$' (compound 10) (224.5 mg. 0.25 mmoles) and diisopropylethylamine (DIPEA) (130.64 µl, 0.75 mmoles) were added at rt. The whole reaction mixture was allowed to stir under argon at 25°C for 36 hrs, then treated with ether: petroleum ether 1:1 (100 ml). The precipitate was filtered, washed with ether, and air dried to afford an off-white solid (197 mg), which was then subjected to HPLC separation and purification.

Analytical HPLC: Pep1 gradient; Gemini C18 column 100 A 5 µm 150 x 3.00 mm; wavelength 220/280 nm; flow rate 0.7 ml/min; solvent A = 0.1% Trifluoroacetic acid, solvent B = 100% acetonitrile; temperature 30°C; gradient: 0-50% B, 15 min. Retention time: compound 12 at 14.35 min, compound 13 at 14.53 min.

HPLC preparative: Water Xterra C18 prep MS Column 19 x 50 mm, 5 µm. Wavelength 220/280nm. Flow rate 8 ml/min. Solvent A = 0.1% Trifluoroacetic acid. Solvent B = 100% acetonitrile. Temperature 30°C. Gradient: 10-100% B, 80 min. Pure fractions of compounds 12 and 13 were collected and freeze dried separately to afford PKI PYR($Z_n$')2 (compound 12) 49 mg, and PK2PYR($Z_n$')2 (compound 13) 42 mg, respectively.

PK1PYR(Zn')2 (compound 12) MS indicated ESI +ve 10V 1331.5 ion. 1H-NMR ($D_20$) δ (ppm): 8.00 (2H, s, aromatic para H x 2), 6.00 (2H, d, H-3 x 2), 5.20-5.50 (4H, m, H-7 x 2, H-8 x 2), 4.35-4.90 (8H, m, H-4 x 2, H-5 x 2, H-6 x 2, H-9 x 2), 4.15 (2H, dd, H-9' x 2), 3.80 (6H, s, $COOCH_3$ x 2), 3.25-3.45 (4H, m, $OCONHCH_2$ x 2), 2.90-3.20 (4H, m, $NHCH_2$ x 2), 1.95 (6H, s, $CH_3C0$ x 2), 1.20-1.70 (28H, m, $(CH_2)_7$ x 2).

[0245] PK2PYR($Z_n$')2 (compound 13) MS indicated ESI +ve 10V 1331.5 ion. 1H-NMR ($D_20$) δ (ppm) 8.35 (1H, s, aromatic H x 1), 7.50 (1H, s, aromatic H x 1), 6.00 (2H, d, H-3 x 2), 5.20-5.50 (4H, m, H-7 x 2, H-8 x 2), 4.35-4.90 (8H, m, H-4 x 2, H-5 x 2, H-6 x 2, H-9 x 2), 4.15 (2H, dd, H-9' x 2), 3.80 (6H, s, $COOCH_3$ x 2),3.25-3.45 (4H, m, $OCONHCH_2$ x 2), 2.90-3.20 (4H, m, $NHCH_2$ x 2), 1.95 (6H, $CH_3CO$ x 2), 1.20-1.70 (28H, m, $(CH_2)_7$ x 2).

**Synthesis of MD185 (PK2 PYR(Zn)2[D-Asp(NHCH2S03H)2]2)**

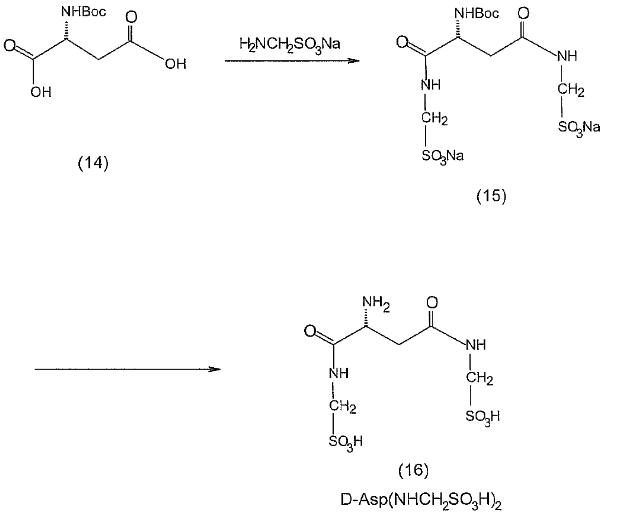


Boc-D-Aspartic acid (14) (233 mg, 1 mmole) and HBTU [O-(benzotriazol-l-yl)- N,N,N',N'-bis(tetramethylene)uronium hexafluorophosphate] (758 mg, 2 mmoles) were dissolved in DMF (10 ml). To this solution DIPEA (diisopropylethylamine) (349µ1, 2 mmoles) was added. The solution was stirred at rt for 10 min, then combined with a solution of sodium aminomethanesulfonate (332 mg, 2.5 mmoles) in DMF (3 ml). The resulting solution was allowed to stir at rt overnight. The reaction mixture was vacuum evaporated to afford a pale orange solid. It was dissolved in water (20 ml), washed with ethyl acetate (20 ml x 3), the aqueous phase was vacuum evaporated to remove organic solvent, then freeze dried to afford crude product (compound 15) 310 mg as a white solid which was subjected to HPLC separation and purification.

HPLC analytical: Pep1gradient; Gemini C18 column 100 A 5 µm 150 x 30 min; wavelength 220 nm; flow rate 0.7 ml/min; solvent A = 0.1% Trifluoroacetic acid, solvent B = 100% acetonitrile; temperature 30°C; gradient 0-50% B, 15 min. Retention time: Compound (15) as free acid at 4.669 min MS ESI -ve 418(M-1), solvent peak at 1.658 min, aminomethanesulfonic acid at 2.722 min, HBTU substance at 5.520 min, mono B0C-D-ASP-$NHCH_2SO_3H$ at 5.802 min.

HPLC preparative: Gemini AXIA packed C18 column, 5 m 50 x 21.2 mm Wavelength 220 min. Flow rate 8 ml/min; Solvent A = 0.1% Trifluoroacetic acid; Solvent B = 100% acetonitrile; Temperature: 30°C. Gradient : 0-100% B, 100 min.

Fractions containing compound 15 were collected and pooled together, then freeze dried to afford pure compound 15 as a free acid. MS ESI -ve 418(M-1). The free acid (200 mg. 0.477 mmoles) was dissolved in trifluoroacetic acid (10 ml) containing Anisole (1 ml). The solution was stirred at rt for 1 hr, then vacuum evaporated to dryness. The residue was triturated in ether (20 ml x 2) and filtered off. The solid was air dried, then redissolved in water, and freeze dried again to afford compound 16 (106 mg) as a white solid. MS ESI -ve 318(M-1). The product was dissolved in water (10 ml), then neutralized with two equivalent of sodium hydroxide, freeze dried to give the disodium salt of compound 16 as a white solid.

Synthesis of MD185 proceeded as in the figure below.

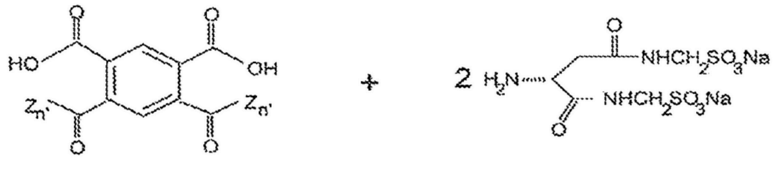


i 16) as di Na salt

(13)

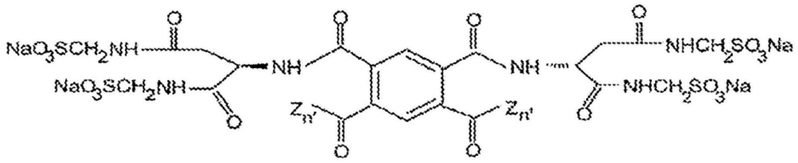


(17)

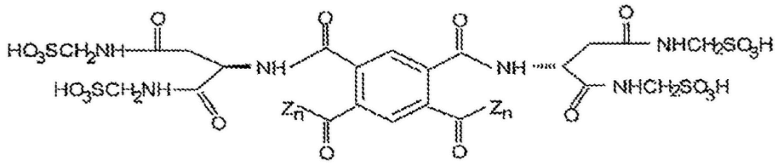


(18)

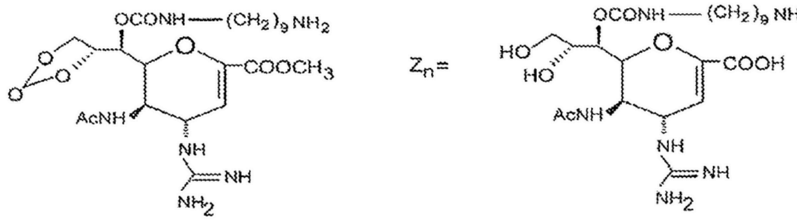


To a solution of 30 mg PK2 PYR(Zn)2 (compound 13) in 5 ml DMF, 17.14 mg HATU [0-(7-Aza benzotriazol-l-yl )-N,N,N',N '-bis(tetramethylene)uranium hexafluorophosphate] and 7.9 µl DIPEA were added. The resulting solution was stirred at room temperature for 10 min, followed by the addition of 33 mg disodium salt of D-Asp(NHCH2S03Na)2 (compound 16) in 5 ml DMF. The reaction mixture was stirred at rt for 18 hrs, then it was subjected to HPLC separation and purification.

HPLC analytical: pep 1 gradient. Gemini C18 column 100A 5µm 150 x 3.00 mm; Wavelength: 220/280 nm; Flow rate : 0.7 ml/min; Solvent A = 0.1% Trifluoroacetic acid; Solvent B = 100% Acetonitrile; Temperature: 30°C; Gradient: 0-50% B, 15 min; Retention time: The product (17) as free acid at 14.270 min. MS ESI +ve +30V

967.5, MS ESI -ve, -25v 965.4. The other peaks were as follows: the starting material PK2PYR($Z_n$') (13) at 11.7 min, PK2PYR($Z_n$')2-D-Asp($NHCH_2S0_3H$)2 at 13.6 min.

HPLC preparative: Water Xterra prep MS C18 column 19 x 50 mm 5 µm; Wavelength: 220/280 nm; Flow rate: 8 ml/min; Solvent A = 0.1% Trifluoroacetic acid; Solvent B = 100% Acetonitrile; Temperature: 30°C; Gradient : 0-100%B 100 min

Product elutes in reverse profile to Gemini analytical. Elution profile between 25-35% acetonitrile. Fractions sampled and tubes with greater than 90% purity were pooled and other material was separated on the Gemini AXIA column. Product (17) was freeze dried to afford a white fluffy powder. MS ESI -ve -25v 965.4, indicated the MW of 1932.

Compound (17) PK2 PYR($Z_n$ )$_2$[D- Asp($NHCH_2S0_3H$)$_2$]$_2$ '(8 mg, 4.14 µmoles) was dissolved in 50% methanol aqueous solution (2 ml), then triethylamine (40 µ1, 287.5 µmoles) was added The resulting solution was stirred at room temperature. Reaction was monitored by analytical HPLC for peak at 10.848 min. After 2 hrs, the reaction mixture was acidified with acetic acid (60 µl, 1049 µmoles) and diluted to 10 ml with water. The solution was purified on the Kinetex column. Pure fractions were combined and freeze dried to afford product (18) PK2 PYR($Z_n$)$_2$[D-Asp($NHCH_2S0_3H$)$_2$]$_2$ 6 mg as white powder.

HPLC retention time 10.695 min. MS ESI -ve -30V 925.6.

In order to remove residue of TFA in compound, the pure material was dissolved in 60 ml 4 mM HCl and freeze dried, then dissolved in 60 ml 1 mM HCl and freeze dried, and finally freeze dried from 60 ml water. Yield.5.6 mg (18) MD185 MW1852.

**Preparation of Acidic/Anionic Group [D-Cyst-$S0_3H$]$_7$OH**

In the scheme (24) to (31) below, solid phase synthesis was applied to prepare [D-Cys-SH]$_7$-OH (33), then oxidation of the polycysteine (33) provided [D-Cya$S0_3H$]$_7$-OH (34).

Fmoc-D-Cys(Trt)-OH (24) (2.93 g, 5 mmoles) was dissolved in dichloromethane (DCM) (35 ml) in a 50 ml Falcon tube, then was added to 2-chlorotrityl chloride resin (25) (5 g), shaken vigorously, and diisopropylethylamine (DIPEA) (3.5 ml, 20 mmoles) was added. After 5 min, another portion of DIPEA (1.3 ml, 7.5 mmoles) was added. The whole mixture was shaken for a further 4 hrs. To endcap any remaining reactive 2- chlorotrityl chloride, methanol was added (4 ml). The mixture was shaken for further 30 min. The resin was washed successively with DMF (20 ml x 3), DCM (20 ml x 3), methanol (20 ml x 3), then vacuum dried overnight. Yield indicated loading of 0.57 mM/g (26).

For removal of the Fmoc group from the resin, the resin (26) was pre swollen in DCM (35 ml), then 50% piperidine in DMF was added (30 ml). The whole mixture was placed on a rotator for 30 min. This was repeated with a fresh solution of 50% piperidine in DMF (30 ml) for 60 min. Resin was then washed successively with DMF (20 ml x 2), DCM (20 ml x 2), DMF (20 ml x 3). A positive ninhydrin test was presented. The resin (27) was ready for coupling.

To a solution of Fmoc-D-Cys(Trt)-OH (24) (1758 mg, 3 mmoles) in DMF (6 ml) HBTU [0-(Benzotriazol-l-yl)-N,N,N',N'-bis(tetramethylene)uronium hexafluorophosphate] (1140 mg, 3 mmoles) and DIPEA (523 ul, 3 mmoles) were added. The mixture was stirred at room temperature for 10 min, then resin was added (27) (2.5 g, 0.57 mM/g, 1.5 mmoles). The reaction mixture was allowed to agitate for 24 hrs. Ninhydrin test negative, the resin was washed with DMF (20 ml x 3), then treated with 5% acetic anhydride, 1% DIPEA in DMF (20 ml) for 30 min. The resin (28) was then washed with DMF (20 ml x 3), DCM (20 ml x 3) and methanol (20 ml x 3). The resin (28) was pre swollen in DCM, and 30% piperidine/N-methylpyrollidone in DMF (20 ml) was added. The whole mixture was placed on a rotator for 30 min, then with a fresh solution of 30% piperidine and N-methylpyrollidone for a further 60 min, after washing with DMF (20 ml x 2), DCM (20 ml x 2), DMF (20 ml x 2) to afford resin (29).

The resin (29) was further coupled for 24 hrs. with (24) (1758 mg, 3 mmoles) in DMF (6 ml) preactivated for 10 min with HBTU (1140 mg, 3 mmoles) and DIPEA (523 uµl, 3 mmoles), after work-up to afford resin (30). Half of which was then: deprotected by piperidine/methylpyrollidone, recoupled with (24) preactivated with HBTU/DIPEA, and this procedure was repeated 4 times to produce resin (31).

Resin (31) was washed with DCM (20 ml x 5), then was placed in a 50 ml Falcon tube with 40% acetic acid in DCM (35 ml), placed on a rotator at room temperature and shaken for 6 hrs. The resin suspension was filtered, the filtrate was vacuum evaporated to a pale yellow foam. The residue was washed with water (100 ml x 3), then dried to afford a white solid (32) $[D\text{-}Cys(Trt)]_7$-OH.

(32) was stirred in a solution of TFA (20 ml), Triisopropylsilane (1 ml) and water (0.5 ml) for 3 hrs. The resulting suspension was filtered. The filtrate was evaporated to an oily substance which was then triturated with ether (20 ml x 4) to give a white solid. It was stirred in 50% acetonitrile aqueous solution (100 ml). The material was sonicated to form a white suspension, which was freeze dried to afford a crude product of (33) $[D\text{-}Cys\text{-} SH]_7$-OH (380 mg).

> HPLC analytical: Phenomenex Gemini 5 µm C18 110A column 150 x 3.00 mm Solvent A = 0.1% Trifluoroacetic acid, Solvent B= 100% Acetonitrile, Flow rate: 0.7 ml/min, Wavelength: 210/280 nm, Temperature: 30°C, Gradient: 0-50% B 15 min, Retention time: 9.356 min.
>
> MS Cone +25 V Major peak 740
> MS Cone -25V Major peak 738
> Indicated MW of (33) is 739 $[D\text{-}Cys\text{-}SH]_7$-OH

Compound (33) (10 mg, 13.53 µmoles) was added to a performic acid solution* (1 ml) prechilled in an ice-salt bath. The whole mixture was stirred in the ice bath for 1 hr, then diluted with 100 ml cold water, freeze dried to afford a pale yellow form (34). This material was purified on a Phenomenex Kinetex 5 µm XB-C18 100A column using isocratic 0.1% TFA as eluent to afford compound (34) $[D\text{-}Cyst\text{-}S0_3H]_7$-OH MW 1075, MS Cone +50V 1076 (M+1) as a colourless powder.:

*Performic acid was prepared by mixing Formic acid (8.74 ml), water (0.96 ml) and 30% hydrogen peroxide (1 ml) and allowing the mixture to stand at room temperature for 30 min in a stopped flask. This peroxide should be freshly made before use.

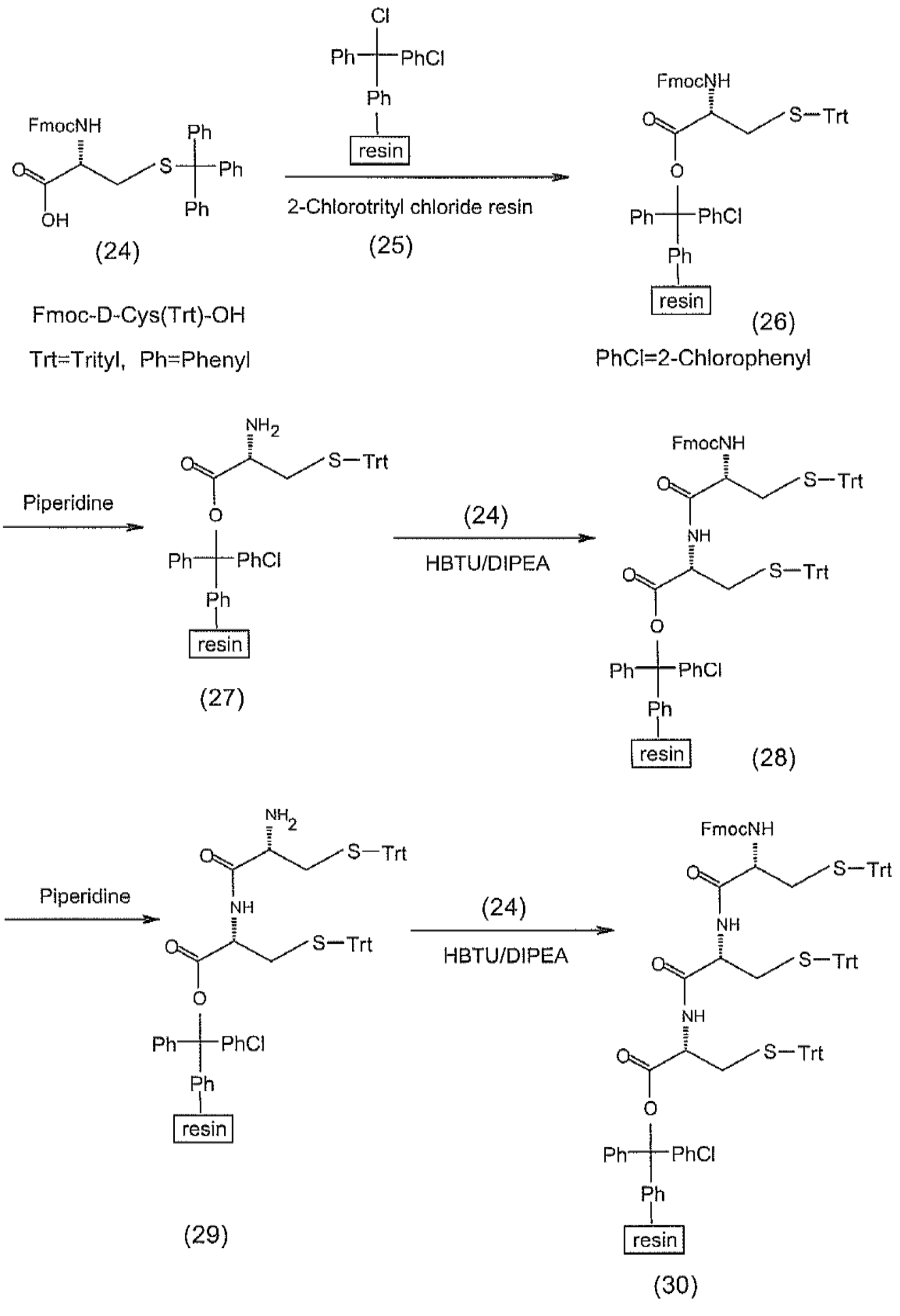
Cl
Ph
PhCl
Ph
resin
FmocNH
Ph
S
Ph
O
OH
Ph
(24)
2-Chlorotrityl chloride resin
(25)
FmocNH
S—Trt
O
O
Ph
PhCl
Ph
resin
(26)
Fmoc-D-Cys(Trt)-OH
Trt=Trityl, Ph=Phenyl
PhCl=2-Chlorophenyl
Piperidine
NH2
S—Trt
O
O
Ph
PhCl
Ph
resin
(27)
(24)
HBTU/DIPEA
FmocNH
S—Trt
O
NH
S—Trt
O
O
Ph
PhCl
Ph
resin
(28)
Piperidine
NH2
S—Trt
O
NH
S—Trt
O
O
Ph
PhCl
Ph
resin
(29)
(24)
HBTU/DIPEA
FmocNH
S—Trt
O
NH
S—Trt
O
NH
S—Trt
O
O
Ph
PhCl
Ph
resin
(30)

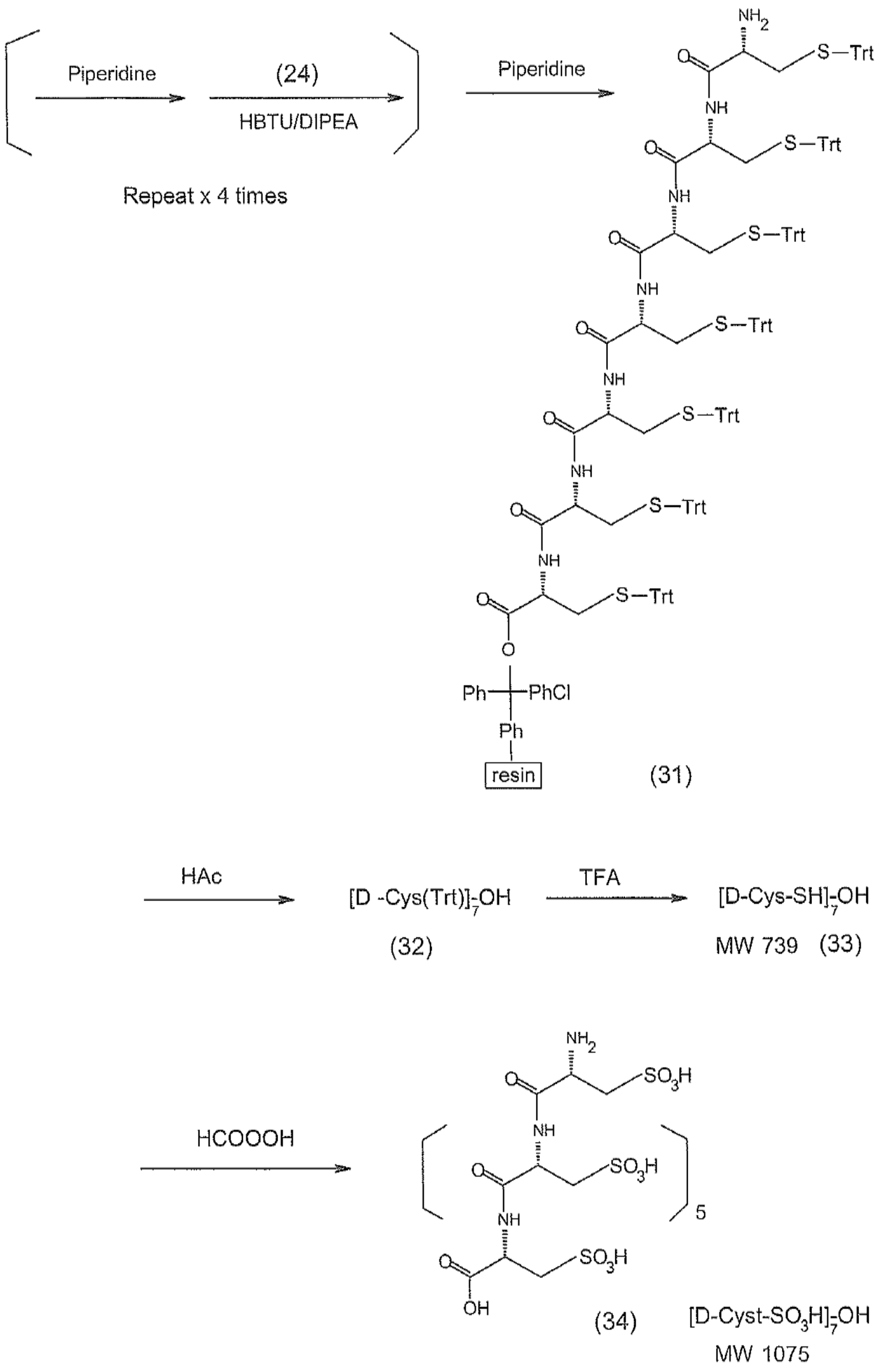

Piperidine
(24)
HBTU/DIPEA
Repeat x 4 times
Piperidine
NH2
S—Trt
O
NH
Ph
PhCl
resin
(31)
HAc
[D -Cys(Trt)]7-OH
(32)
TFA
[D-Cys-SH]7-OH
MW 739
(33)
HCOOOH
SO3H
5
OH
(34)
[D-Cyst-SO3H]7-OH
MW 1075

**Preparation of Compound MD345, PK2 TCA(Zo)2-[D-Cyst-S03H]7-OH MW2201**

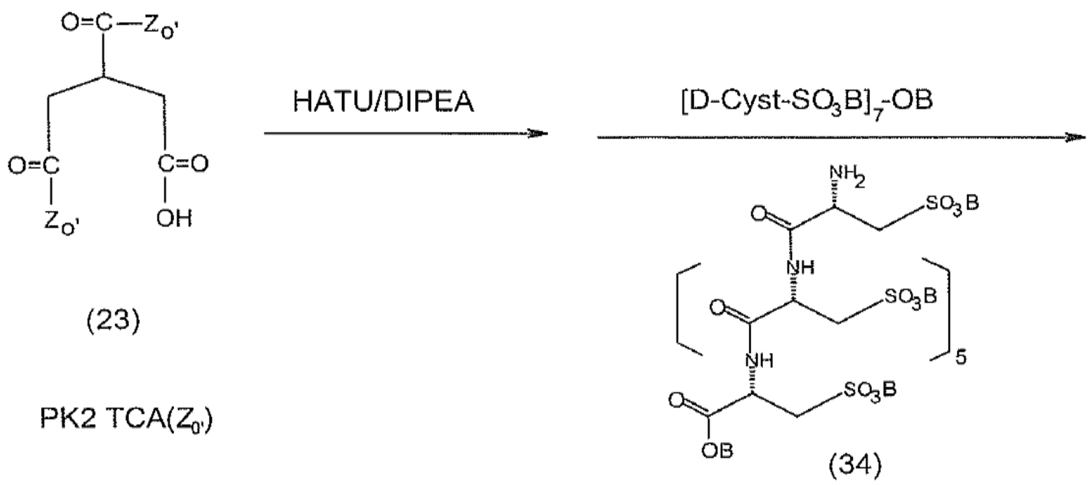


(23)

PK2 TCA($Z_{0'}$)

(34)

B=base, DIPEA

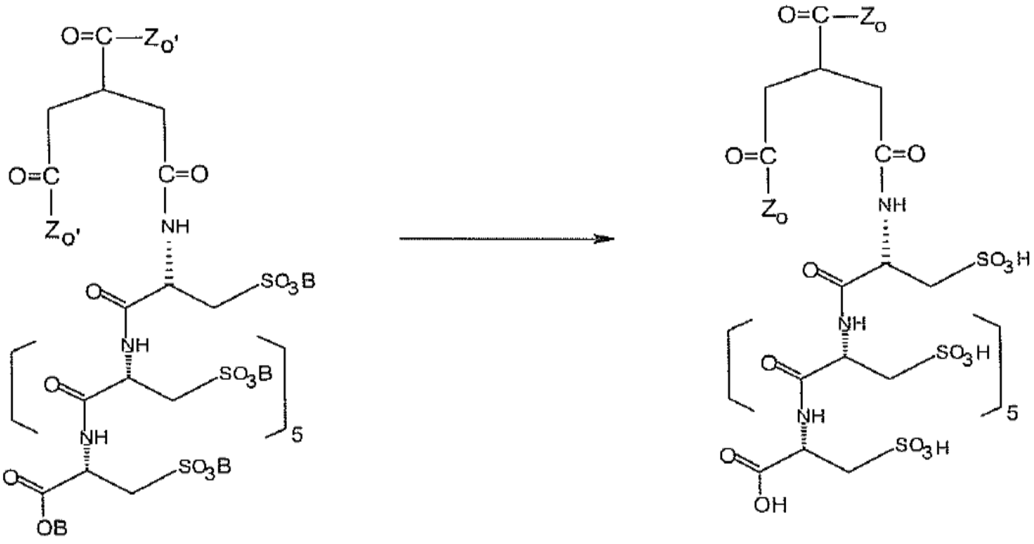

(35)

PK2 TCA($Z_{0'}$)$_2$-[D-Cyst-So$_3$B]$_7$-OB

(36)

PK2 TCA($z_0$)$_2$-[D-Cyst-SO$_3$H]$_7$-OH

$Z_{0'}$ =

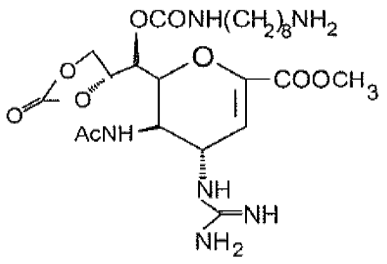

$Z_0$ =

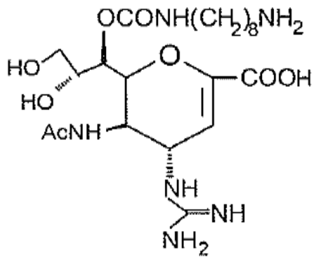

To a solution of PK2 TCA($Z_0$')$_2$ (23) (4 mg, 3.268 µmoles) in anhydrous DMF (2 ml) HATU (1.242 mg, 3.268 µmoles) and 2% DIPEA in DMF (30 µl, 3.44 µmoles) were added. The mixture was stirred at room temperature for 10 min, then combined with a solution of (34), [D-Cyst-$S0_3$H]$_7$-OH (7.03 mg, 6.54 µmoles) and DIPEA (9.11 µl, 52.32 µmoles), in DMF (3 ml). The whole reaction mixture was stirred at room temperature overnight. The mixture was diluted to 15 ml with water/methanol 1:1, then subjected to HPLC separation and purification.

> HPLC preparative: Phenomenex Kinetex 5 µm XB-C18 column 50 x 21.2 mm; Wavelength: 220/280 nm; Flow rate: 8 ml/min; Solvent A = 0.1% Trifluoroacetic acid (TFA); Solvent B = 100% Acetonitrile; Temperature: 30°C; Gradient: 0-100% B, 80 minutes.

Fractions containing (35) free acid (B=H) were combined and freeze dried to afford compound (35) free acid (B=H) 4 mg as a white powder. MS ESI +ve Cone 35V 1142.1 2 + ion.

Compound (35) free acid (B=H), PK2 TCA($Z_0$')$_2$-[D-Cyst-$SO_3$H]$_7$-OH (4 mg, 1.754 µmoles) was dissolved in 50% methanol aqueous solution (800 µl) containing triethylamine (TEA) (20 µl, 146 µmoles). The mixture was stirred at room temperature, monitored by HPLC. It was worked up after 3 hrs. The solution was purified on the Kinetex column. Pure fractions at a retention time of 11.254 min were combined and freeze dried to afford compound (36) 2 mg as a white powder. MS ESI +ve Cone 40V 1102.2 2 + ion.

In order to remove the residue of TFA in the compound, compound (36) was dissolved in 4 mM HCl (30 ml) and freeze dried. Then it was dissolved in 1 mM HCl (30 ml) and freeze dried. Finally it was freeze dried from water (30 ml) to afford product (36) 1.1 mg, MD345 PK2 TCA($Z_0$)$_2$[D-Cyst-$SO_3$H]$_7$-OH MW2201 [MS 2202 (M+1)].
