## Supplementary Figure 1 for "A new concept in antiviral drug design yields a potent influenza inhibitor"

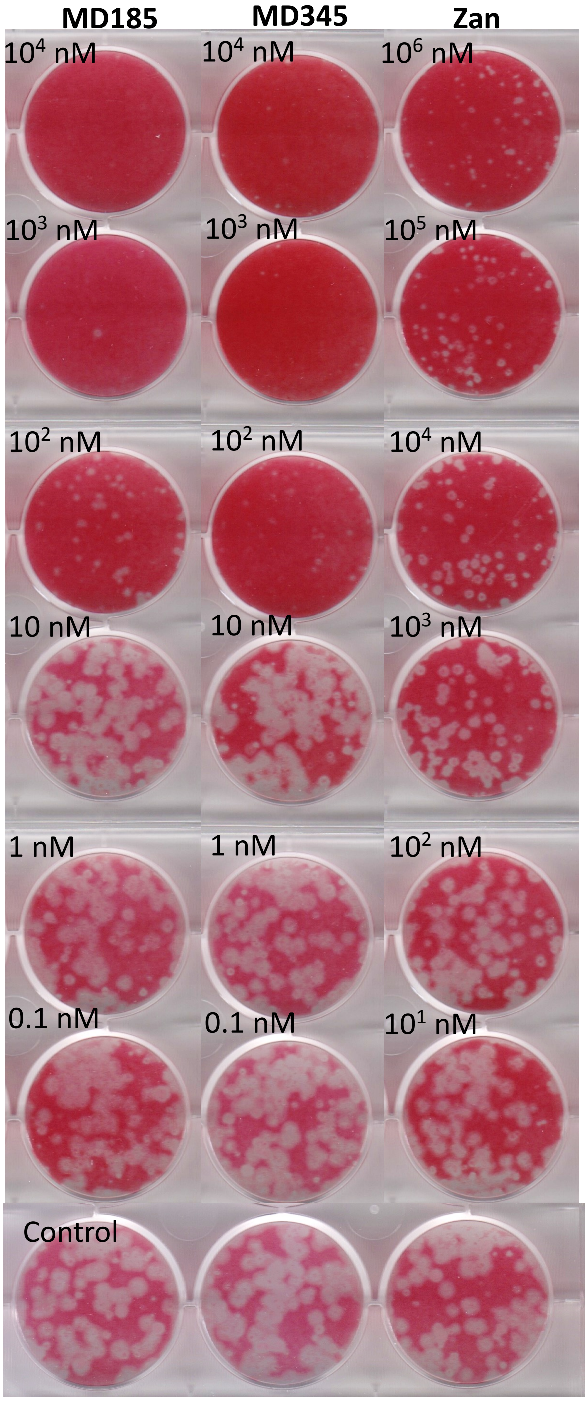


**Supplementary Figure 1.** Plaque assay in the presence of the compounds showing reduction in number of plaques as the primary outcome with MD185 and MD345 compared to zanamivir where reduction in plaque size was the predominant result. Combined images of wells from 6-well plates containing plaque assays of A/Switzerland/9715293/2013 virus in MDCK cells in which the virus was mixed with MD185, MD345 or zanamivir at the concentrations shown before addition to cells. Monolayers were stained with neutral red. Note the difference in the concentration series of zanamivir compared to the other two compounds.
