## Supplementary Figure 2 for "A new concept in antiviral drug design yields a potent influenza inhibitor"

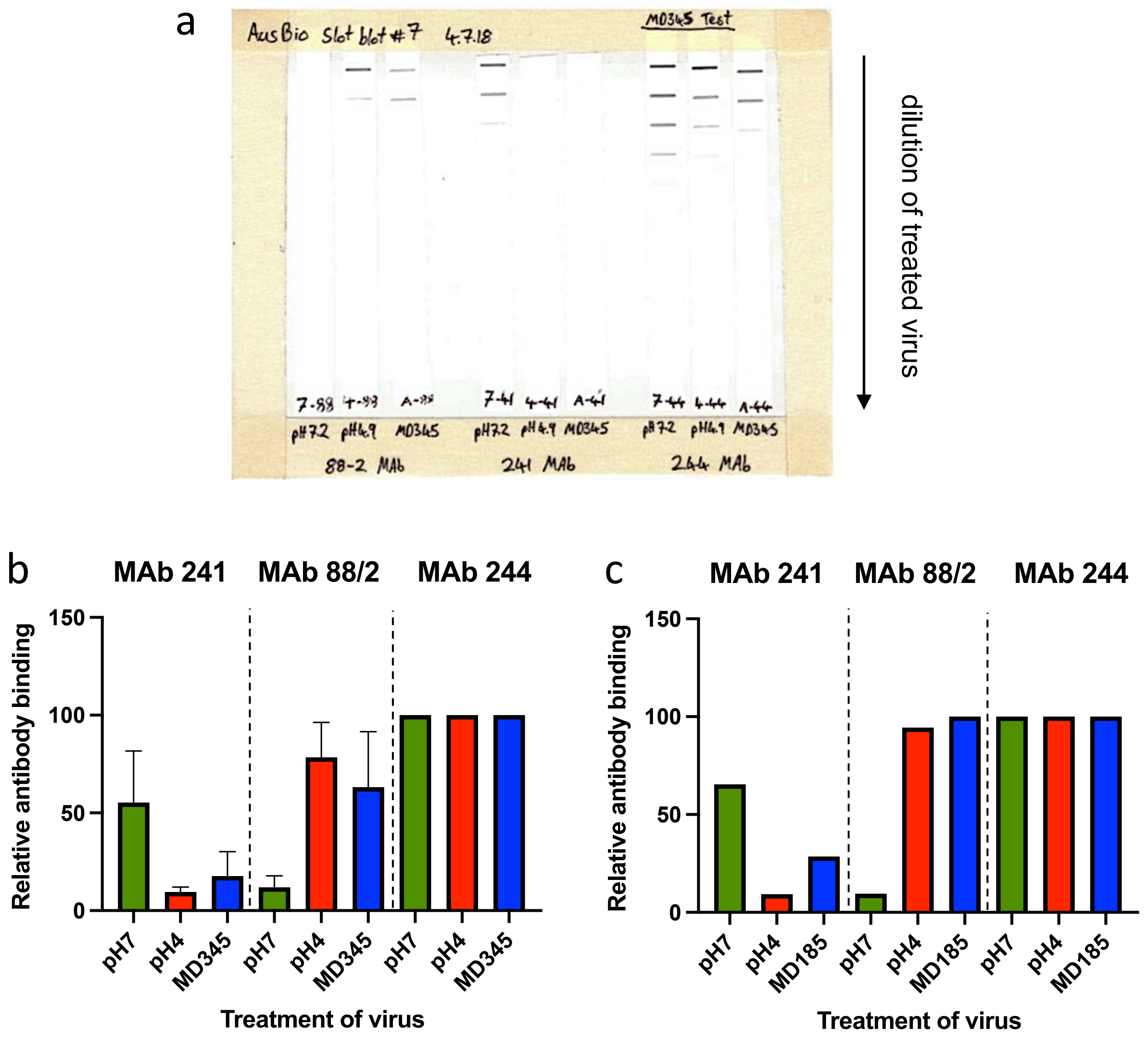


**Supplementary Figure 2.** Slot blot assay. Panel **a** shows the stained nitrocellulose strips used to calculate the relative binding of MAb 241 (specific for the neutral pH conformation of HA), 88/2 (specific for the low pH conformation of HA) and MAb 244 (binds to both conformations) to pH 7.2-, pH 4.9- or MD345-treated A/Memphis/1/71 virus used for the example in Fig. 2a. of the main paper. Panel **b** shows the relative binding of the MAbs averaged over 3 independent experiments where A/Memphis/1/71-Bel/42 virus ($3.4x10^7$ PFU/mL) was treated with an excess of MD345 (180-360 µg) for 2 hr before blotting and staining. The staining levels for each treatment are standardized to the staining obtained with MAb 244. Data for MD345 treatment was not significantly different to that of the low pH treatment, one-way ANOVA. The pattern of reactivity is analogous to that observed in and additional experiment treating the virus with MD185 (700 µg) (**c**).
