## Supplementary Figure 3 for "A new concept in antiviral drug design yields a potent influenza inhibitor"

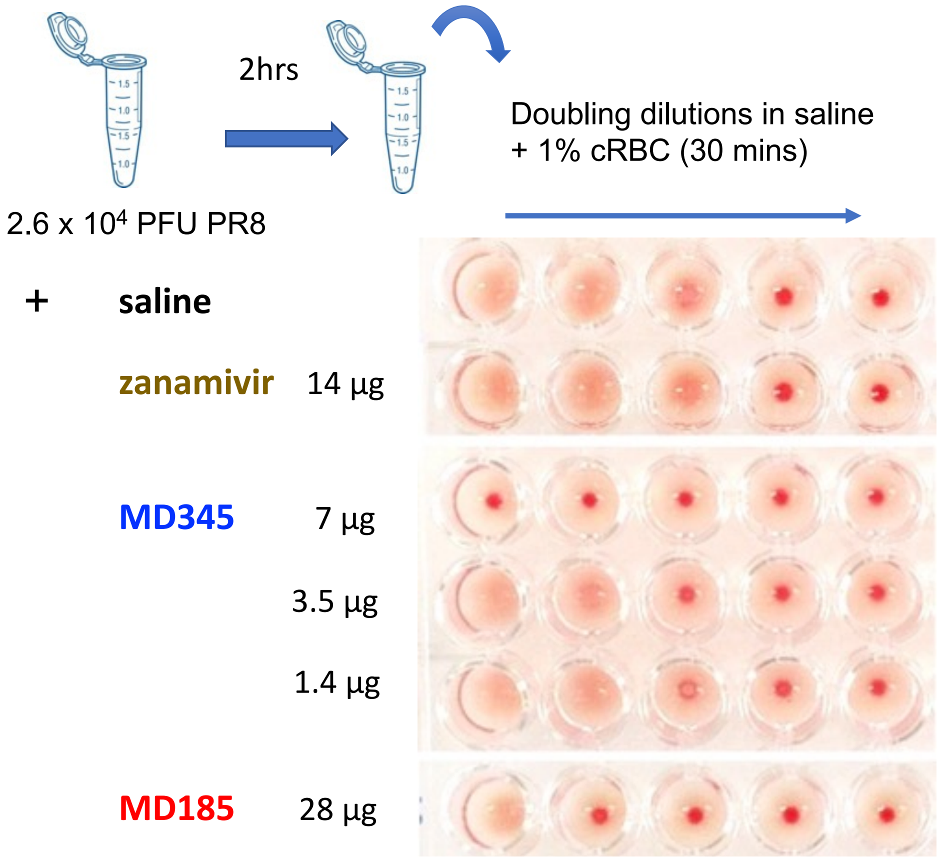


**Supplementary Figure 3.** Inhibition of hemagglutination by MD345 and MD185. PR8 virus was mixed with saline or the compounds as shown in the schematic and after 2 h an HA assay was performed. With saline, this amount of virus caused hemagglutination in the first 3 wells of the dilution series and this pattern was no different in the presence of zanamivir. In contrast, MD345 and MD185 were able to inhibit the virus binding to red blood cells via its HA. MD345 was superior to MD185 in that complete inhibition was observed in the presence of 7 µg of MD345, while it took 28 µg of MD185 to reduce the hemagglutination to the first well. These results represent experiments performed at least 3 times.
