## Supplementary Figure 4 for "A new concept in antiviral drug design yields a potent influenza inhibitor"

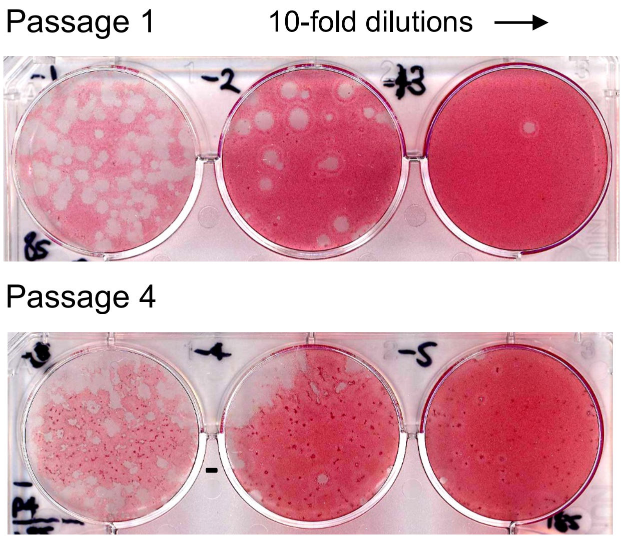


**Supplementary Figure 4.** Plaque morphology of wild type and mutant virus populations. A/Perth/265/09 (pdmH1N1) virus was passaged in MDCK cells in the presence of 10 ng/ml of MD185. After four passages the morphology of the virus population had changed when titrated in a plaque assay in the absence of compound. Images shown represent monolayers stained with neutral red depicting the virus morphology after the first passage, which comprised predominantly medium to large clear plaques, compared to the fourth passage, which showed a minority of larger clear plaques that proved to be wild type virus when sequenced, and a predominance of very small plaques with a dense center surrounded by a semi-transparent ring. Two of the smaller plaques were amplified and found to have NA Q136K or NA E119K + HA G172E substitutions.
