## Supplementary Figure 5 for "A new concept in antiviral drug design yields a potent influenza inhibitor"

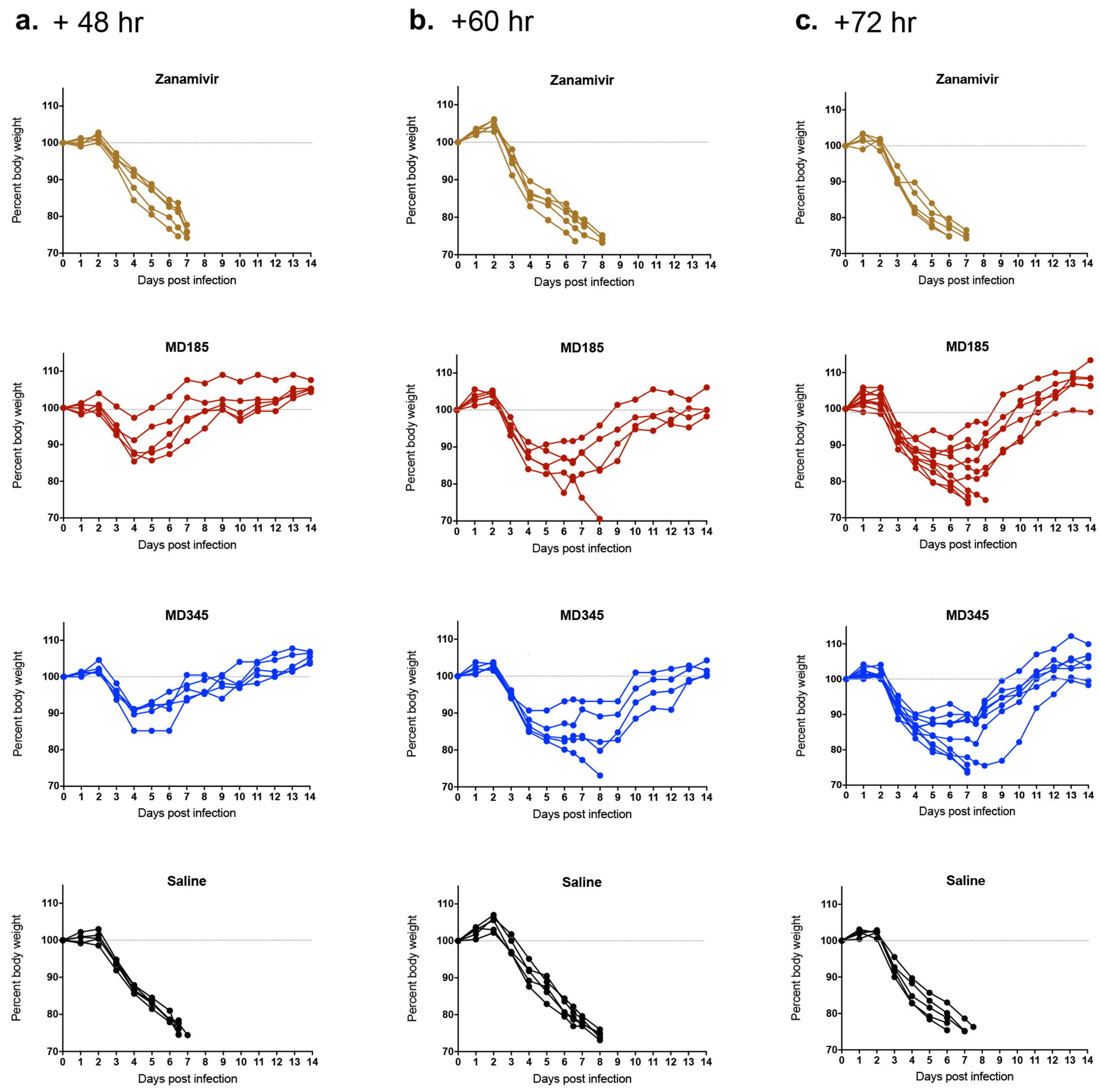


**Supplementary Figure 5.** Individual mouse weights after infection and treatment with compound. Mice (5 per group, or 10 per group for the compounds at 72 h) were infected with a lethal dose of PR8 virus followed by treatment with a single 40 µg dose of either zanamivir, MD185, MD345 or with saline at 48 h (column **a.**), 60 h (column **b.**), or 72 h (column **c.**) post infection. Weights and clinical signs were monitored at least once daily and mice were euthanised at a predetermined humane endpoint if required. Graphs show weights of individual mice, expressed as a percentage of starting weight, that are summarised in Fig. 4a of the main paper.
