## Supplementary Figure 6 for "A new concept in antiviral drug design yields a potent influenza inhibitor"

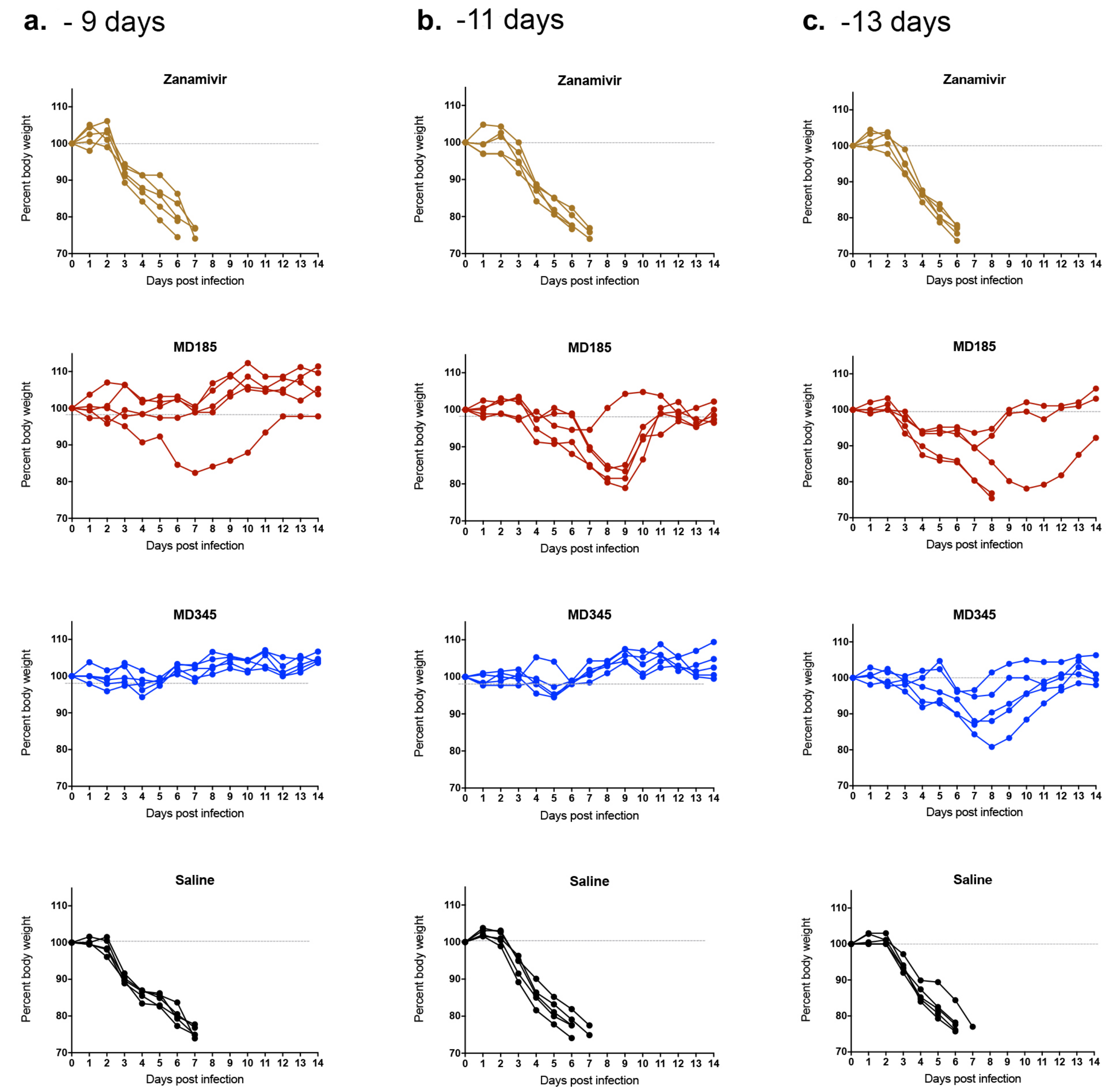


**Supplementary Figure 6.** Individual mouse weights after prophylaxis with compound and infection with virus. Mice (5 per group) were given a single 5 µg dose of either zanamivir, MD185 or MD345, or saline at 9 days (column **a**.), 11 days (column **b**.), or 13 days (column **c**.) before infection with a lethal dose of PR8 virus. Weights and clinical signs were monitored at least once daily and mice were euthanised at a predetermined humane endpoint if required. Graphs show weights of individual mice, expressed as a percentage of starting weight, that are summarised in Fig. 4b of the main paper.
