## Supplementary Figure 7 for "A new concept in antiviral drug design yields a potent influenza inhibitor"

**a**

| Donor Ferret # | $Log_{10}$ $TCID_{50}$/ mL |
|---|---|
| 1 | 3.83 |
| 2 | 3.83 |
| 3 | 4.50 |
| 4 | 3.83 |
| 5 | 3.83 |
| 6 | 3.17 |
| 7 | 3.50 |
| 8 | 4.50 |
| 9 | 3.50 |
| 10 | 3.50 |

**b**

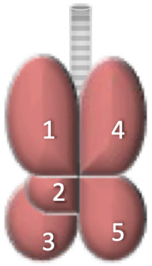


**c**

| | | Pulmonary virus titre ($Log_{10}$ $TCID_{50}$/ml) | | | | |
|---|---|---|---|---|---|---|
| | | Lung lobe | | | | |
| Ferret # | Group | 1 | 2 | 3 | 4 | 5 |
| 11 | 1: MD185-1x | - | - | - | - | - |
| 12 | | - | - | - | - | - |
| 13 | | - | - | - | - | - |
| 14 | | - | - | - | - | - |
| 15 | 2: MD185-5x | - | - | - | - | - |
| 16 | | - | - | - | - | - |
| 17 | | - | - | - | - | - |
| 18 | | - | - | - | - | - |
| 19 | 3: Zanamivir | - | 4.50 | - | 3.17 | - |
| 20 | | 3.00 | - | 3.50 | - | - |
| 21 | | - | - | - | - | - |
| 22 | | 2.50 | 4.50 | - | 2.50 | - |
| 23 | 4: Placebo | - | - | 2.50 | 3.50 | - |
| 24 | | 3.17 | 2.50 | 2.83 | 2.83 | - |
| 25 | | 4.83 | 4.83 | 3.50 | 5.17 | 4.17 |
| 26 | | - | 4.83 | - | - | 3.00 |
| 27 | 5: Oseltamivir | - | - | - | - | - |
| 28 | | - | - | - | - | - |
| 29 | | - | - | - | - | - |

**d**

| | | Infectious virus in nasal wash ($Log_{10}$ $TCID_{50}$/ml) | | | | | | |
|---|---|---|---|---|---|---|---|---|
| | | Day post infection | | | | | | |
| Ferret # | Group | 1 | 2 | 3 | 4 | 5 | 6 | 7 |
| 11 | 1: MD185-1x | No nasal wash | | | | | | - |
| 12 | | | | | | | | - |
| 13 | | | | | | | | - |
| 14 | | | | | | | | - |
| 15 | 2: MD185-5x | - | - | - | - | - | - | - |
| 16 | | - | - | - | - | - | - | - |
| 17 | | - | - | - | - | - | - | - |
| 18 | | - | - | - | - | - | - | - |
| 19 | 3: Zanamivir | - | - | - | - | - | - | 3.17 |
| 20 | | - | - | - | - | - | 2.50 | 2.83 |
| 21 | | - | - | - | - | - | - | - |
| 22 | | - | - | - | - | - | - | - |
| 23 | 4: Placebo | - | - | - | - | - | - | - |
| 24 | | - | - | 4.50 | 5.83 | 4.50 | 3.50 | 3.17 |
| 25 | | - | - | - | 3.83 | 5.17 | 4.50 | - |
| 26 | | - | - | - | - | - | 3.17 | 6.50 |
| 27 | 5: Oseltamivir | - | - | - | - | 2.83 | - | 5.17 |
| 28 | | - | - | - | - | - | - | - |
| 29 | | - | - | - | - | - | - | - |
